# β-catenin programs a tissue-specific epigenetic vulnerability in aggressive adrenocortical carcinoma

**DOI:** 10.1101/2022.07.02.497654

**Authors:** Dipika R. Mohan, Kleiton S. Borges, Isabella Finco, Christopher R. LaPensee, Juilee Rege, April L. Solon, Donald W. Little, Tobias Else, Madson Q. Almeida, Derek Dang, James Haggerty-Skeans, April A. Apfelbaum, Michelle Vinco, Alda Wakamatsu, Beatriz MP Mariani, Larissa Amorin, Ana Claudia Latronico, Berenice B. Mendonca, Maria Claudia N. Zerbini, Elizabeth R. Lawlor, Ryoma Ohi, Richard J. Auchus, William E. Rainey, Suely K. N. Marie, Thomas J. Giordano, Sriram Venneti, Maria Candida B. V. Fragoso, David T. Breault, Antonio Marcondes Lerario, Gary D. Hammer

## Abstract

Adrenocortical carcinoma (ACC) is a rare cancer in which tissue-specific differentiation is paradoxically associated with dismal outcomes. The differentiated ACC subtype CIMP-high is prevalent, incurable, and routinely fatal. CIMP-high ACC possess abnormal DNA methylation and frequent β-catenin activating mutations. Here, we demonstrate that ACC differentiation is maintained by a balance between nuclear, tissue-specific β-catenin-containing complexes and the epigenome. On chromatin, β-catenin binds master adrenal transcription factor SF1 and hijacks the adrenocortical super-enhancer landscape to maintain differentiation. Off chromatin, β-catenin binds histone methyltransferase EZH2, which is redistributed by the CIMP-high DNA methylation signature. SF1/β-catenin and EZH2/β-catenin complexes exist in normal adrenals and are selected for through all phases of ACC evolution. Pharmacologic EZH2 inhibition in CIMP-high ACC favors EZH2/β-catenin assembly and purges SF1/β-catenin from chromatin, erasing differentiation and restraining cancer growth *in vitro* and *in vivo*. Our studies illustrate how tissue-specific programs shape oncogene selection, surreptitiously encoding targetable therapeutic vulnerabilities.

## INTRODUCTION

Deranged epigenetic patterning is a hallmark of cancer (Flavahan et al., 2017; Kadoch et al., 2013). CIMP-high is a recurrent epigenetic signature defined by abnormal DNA methylation in promoter CpG islands (CpGi). These CpGi are normally repressed by histone H3 lysine 27 trimethylation (H3K27me3), written by the histone methyltransferase EZH2 as part of the Polycomb repressive complex 2 (PRC2) (Easwaran et al., 2012). PRC2 is required for embryonic and stem cell pluripotency, restricting lineage specification and differentiation (Deevy and Bracken, 2019). EZH2/PRC2 actions are crucial for many cancers (Schuettengruber et al., 2017). Together, these observations have seeded a model that CIMP-high maintains cancer cell stemness, through redundant or cooperative DNA methylation-dependent silencing of PRC2 targets (Baylin and Jones, 2016; Widschwendter et al., 2007).

Though prevalent across cancer types, CIMP-high has variable prognostic significance. In the rare endocrine cancer adrenocortical carcinoma (ACC), the 30-40% of patients with CIMP-high disease experience rapid metastatic recurrence and death (Mohan et al., 2019; Zheng et al., 2016). Genetically, CIMP-high ACC are distinguished by recurrent somatic alterations leading to constitutive activation of the cell cycle and the Wnt/β-catenin pathway (Assie et al., 2014; Mohan *et al*., 2019; Zheng *et al*., 2016). Classically, Wnt pathway activation culminates in β-catenin co-activation of TCF/LEF-driven transcription and expression of programs that facilitate stem cell maintenance (Nusse and Clevers, 2017). This is also true in the adrenal cortex, the tissue of origin of ACC, where cell type specification is established by integration of paracrine and endocrine cues (Lerario et al., 2022). All adrenocortical cells express master transcription factor SF1 (*NR5A1*). Wnt/β-catenin restrains differentiation of SF1+ cells into a population termed the zona fasciculata (zF). This antagonizes pituitary hormone ACTH, which expands and differentiates the zF to produce glucocorticoids (**Figure 1A**).

**Figure 1:**
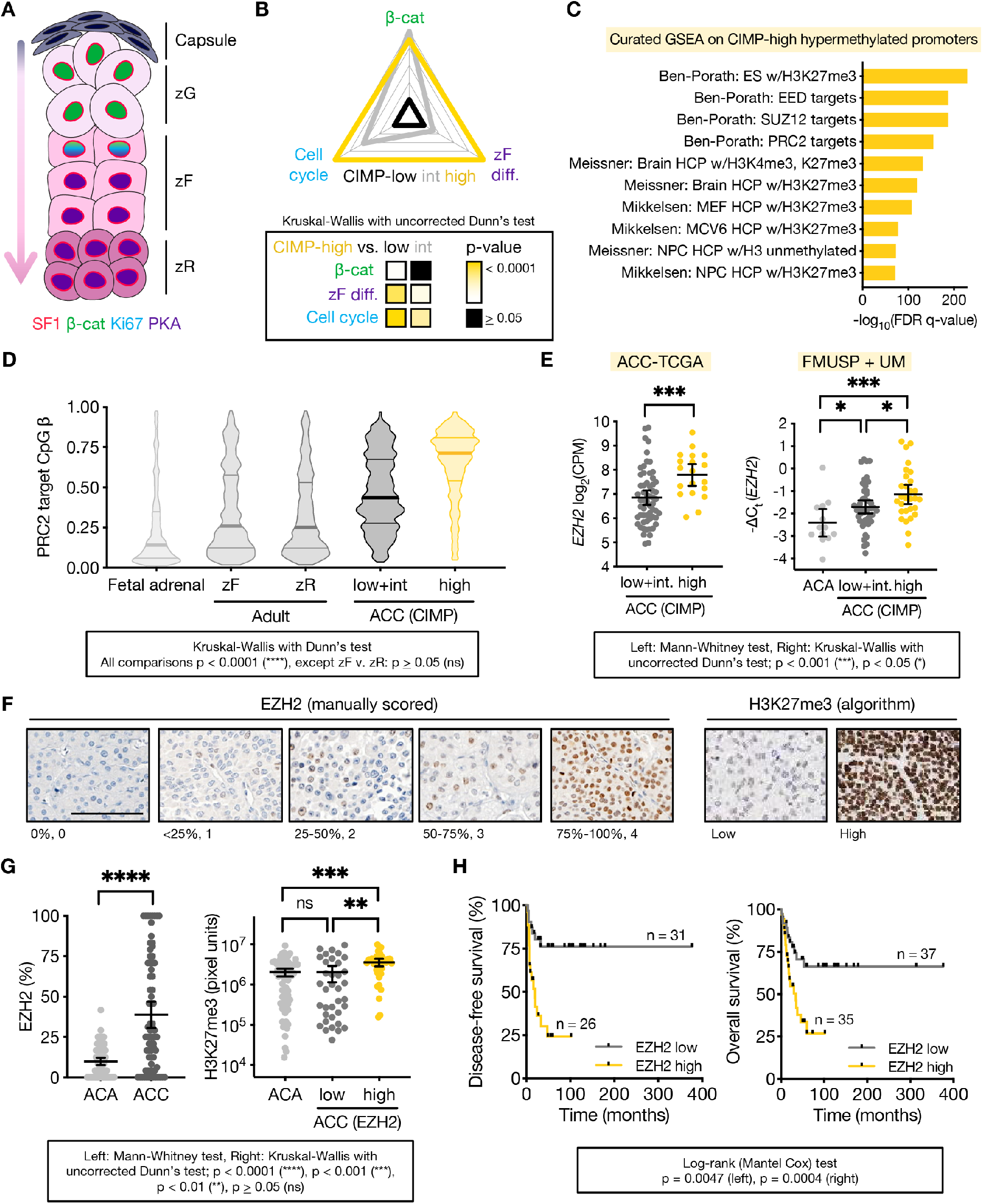
Differentiated, Wnt-active ACC subtype CIMP-high possesses aberrant PRC2 target hypermethylation with high EZH2 coupled to H3K27me3. **(A)** Corticocapsular unit of adrenocortical homeostasis depicting peripheral mesenchymal cells (capsule) and human cortical populations zona glomerulosa (zG), zona fasciculata (zF), and zona reticularis (zR), that produce mineralocorticoids, glucocorticoids, and androgens, respectively. Differentiation in the cortex is centripetal, and zG, zF, and zR cells are supplied by peripheral capsular and cortical progenitors (arrow). The entire cortex is SF1 positive. ZG cells possess active Wnt/β-catenin signaling and lower zF/zR cells possess active ACTH signaling through protein kinase A (PKA). Wnt/β-catenin signaling fades in the upper zF, and these cells respond to ACTH/PKA with proliferation (Ki67). Mice do not have a zR. **(B)** GSVA (Hänzelmann et al., 2013) was used on ACC-TCGA RNA-seq to calculate the expression score of genes that define zF differentiation or are regulated in a cell-cycle- or Wnt-dependent manner across ACC-TCGA. Score validation detailed in **Materials and Methods** and **Supp Fig 1**. Radar plot depicts average score for each ACC CIMP class, with values mapped onto an arbitrary scale along each axis. Heatmap below depicts p-value for comparisons. **(C)** 10 most significant gene sets from curated GSEA (Mootha *et al*., 2003; Subramanian *et al*., 2005) on genes with promoters targeted for hypermethylation in CIMP-high ACC. **(D)** Violin plot of PRC2 target CpG methylation measured by Illumina 850k or 450k methylation array in fetal (n=3) or adult adrenal (zF, zR; n=4 each) and ACC from ACC-TCGA (n=79). Lines at median and quartiles. **(E)** *EZH2* expression in ACC-TCGA by RNA-seq (n=78), left, and independent cohort by qPCR (n=102, FMUSP+UM classified by CIMP in (Mohan *et al*., 2019)), right. CPM=counts per million. ACA=adrenocortical adenomas (benign adrenocortical tumors). Line at mean with 95% confidence interval (CI). **(F-G)** Representative images and scoring of tissue microarray (TMA) of human adult ACA (n=74) and primary ACC (n=74). TMA stained for EZH2 and H3K27me3 by immunohistochemistry (IHC). EZH2 quantified on 0-4 scale (%positive nuclei) by 2 independent observers. H3K27me3 quantified by MATLAB (Bayliss *et al*., 2016). EZH2 low=ACC with below median EZH2 expression (≤25% nuclei EZH2+), EZH2 high=ACC with above median EZH2 expression (>25% nuclei EZH2+). *EZH2* mRNA/protein are correlated (Spearman r=0.5117, p<0.01, not shown). F, bar=100 μm. G, line at mean with 95% CI. **(H)** Disease-free (after R0/RX resection in patients without metastatic disease at diagnosis) and overall survival (all patients) stratified by ACC EZH2 expression.

Broadly, Wnt-dependent tumorigenesis is associated with dedifferentiation, including in CIMP-high neoplasia (Ohm et al., 2007; Schwitalla et al., 2013; Tao et al., 2019; Vaz et al., 2017). Given these observations, one would expect CIMP-high ACC to exhibit dedifferentiation. In striking contrast, across ACC, these cancers exhibit the strongest differentiation, with clinical glucocorticoid production and high expression of zF-defining genes (Mohan *et al*., 2019; Zheng *et al*., 2016). This suggests that aberrant epigenetic programming in ACC stabilizes a paradoxical differentiated, Wnt-active, and rapidly proliferative cellular state (**Figure 1B, Supp Fig 1**). Given advances enabling rapid prospective molecular subtyping of ACC (Mohan *et al*., 2019), the homogeneous clinical and intriguing molecular characteristics of this class, and the abysmal lack of therapies, we investigated the epigenetic underpinnings of CIMP-high ACC.

## RESULTS

### DNA hypermethylation in CIMP-high ACC is pathologically directed to PRC2 targets

DNA hypermethylation is a powerful predictor of survival in ACC; hypermethylation of the *G0S2* CpG island (CpGi) alone captures all features of CIMP-high (Mohan *et al*., 2019). To illuminate mechanisms enabling aberrant epigenetic patterning, we performed GSEA (Mootha et al., 2003; Subramanian et al., 2005) on genes with promoter hypermethylation in CIMP-high tumors from The Cancer Genome Atlas study on ACC (ACC-TCGA; (Zheng *et al*., 2016)). As expected, we observed significant enrichment for embryonic targets of the PRC2 (**Figure 1C**), including *HOX* clusters, *GATA3, PAX6*, and CIMP-high biomarker *G0S2* (**Supp Fig 2A**), mirroring other CIMP-high cancers.

PRC2 targets may gain DNA methylation in mammalian tissues (Chaligne et al., 2021; Tao *et al*., 2019; Toyota et al., 1999; Vaz *et al*., 2017). To determine if CIMP-high methylation reflects tissue origin, we profiled the DNA methylome of fetal and adult adrenals (**Figure 1D**). We observed PRC2 target CpGi acquire minimal methylation during adult differentiation, acquire indiscriminate methylation in non-CIMP-high ACC, and are targeted for methylation in CIMP-high ACC. ACC is devoid of stromal cells ((Zheng *et al*., 2016), **Supp Fig 2B**), suggesting aberrant CpGi methylation originates specifically from cancer cells.

Unlike with *G0S2* (Mohan *et al*., 2019), promoter hypermethylation did not lower gene expression (**Supp Fig 2C**). Given PRC2 targets bear H3K27me3 and exhibit low expression in normal tissue, these data suggest PRC2 target DNA methylation reflects an epigenetic class switch (H3K27me3 exchanged for alternative repressive marks, e.g. DNA methylation and/or H3K9me3 (Gal-Yam et al., 2008; Ohm *et al*., 2007)), or that PRC2 collaborates with DNA methyltransferases (DNMTs) to write DNA methylation at H3K27me3 sites (Viré et al., 2006).

### PRC2 is catalytically active and required for sustained proliferation in CIMP-high ACC

We observed EZH2 is upregulated in a cell-cycle-dependent manner in CIMP-high ACC (**Figure 1E**; **Supp Fig 1E, 2D-F**), as expected (Bracken et al., 2003). EZH2 is nuclear, coupled to high H3K27me3, and predictive of poor clinical outcomes (**Figure 1F-H**). These data suggest EZH2 remains catalytically active on histone substrates in CIMP-high ACC. As EZH2 requires PRC2 incorporation to possess catalytic activity (Cao and Zhang, 2004; Margueron et al., 2009; Montgomery et al., 2005; Pasini et al., 2004), this suggested a major role of EZH2 in ACC involves PRC2.

We then examined if the catalytic function of EZH2 was required for CIMP-high ACC proliferation. Here, we primarily used mainstay human ACC cell line, NCI-H295R, with driver alterations in *TP53, RB1* and *CTNNB1* (encoding β-catenin) (**Supp Fig 3A**). We performed multiplatform profiling of NCI-H295R cells and demonstrated it is a *bona fide* model of CIMP-high ACC (**Figure 2A-C**). We also characterized the murine Y1 and ATC7L ACC cell lines, which harbor genetic alterations in cell cycle machinery characteristic of CIMP-high with variable zF differentiation and Wnt/β-catenin activation (**Supp Fig 3A-G**); however, both lines express *G0s2* (**Supp Fig 3D**).

**Figure 2:**
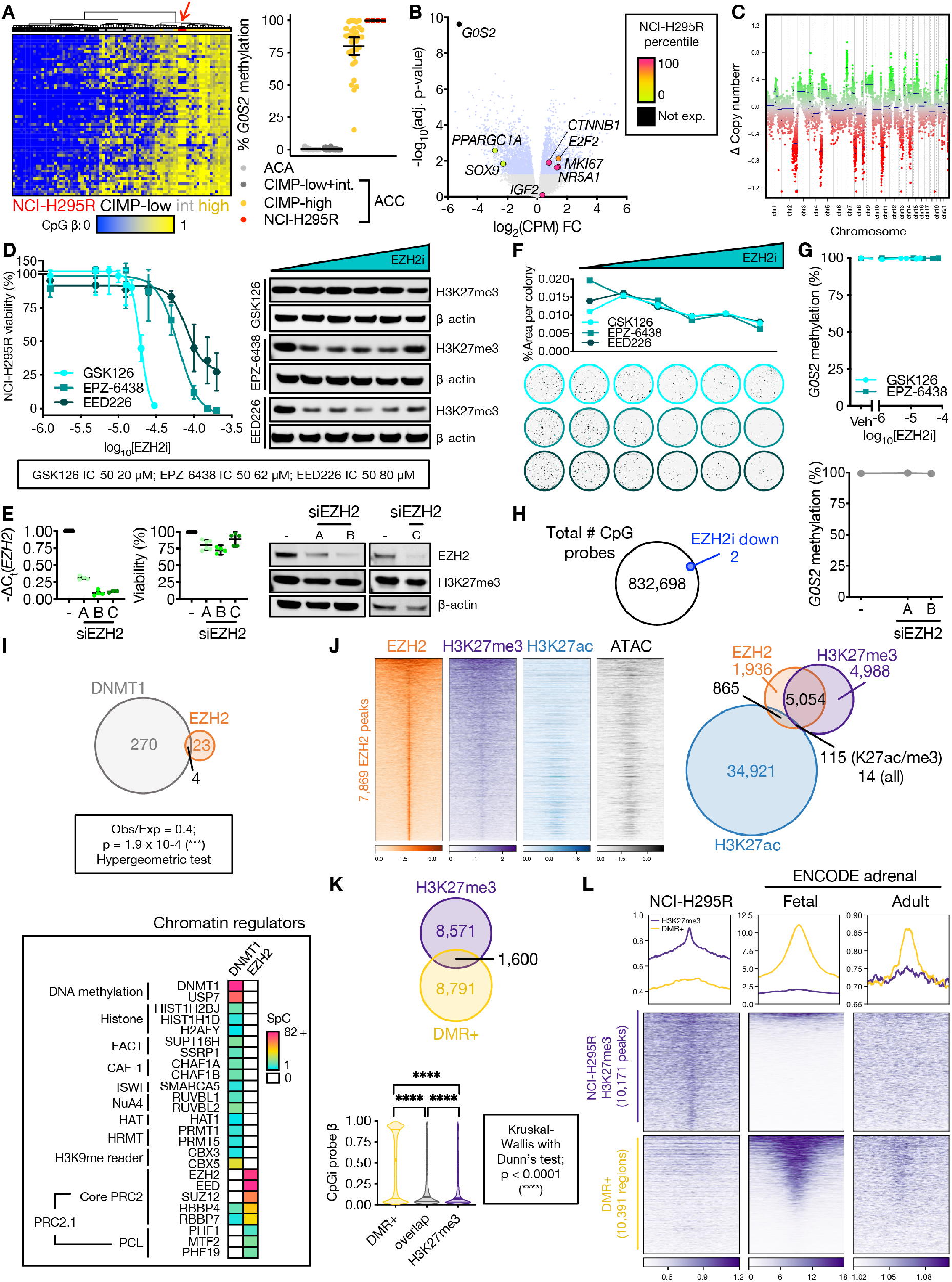
DNA methylation is propagated independently of PRC2 in CIMP-high ACC. **(A)** Left, heatmap of methylation at probes (rows) that define ACC-TCGA CIMP groups. Columns are ACC-TCGA samples classified by CIMP or baseline NCI-H295R cell line (red arrow, n=3, Illumina 850k array). Unsupervised hierarchical clustering with ward.D2 algorithm, Euclidean distance. Right, targeted assessment of *G0S2* methylation in ACA (n=14), ACC stratified by CIMP-status (n=49 non-CIMP-high, n=33 CIMP-high), and baseline NCI-H295R, n=4; line at mean with 95% CI. ACA+ACC data from (Mohan *et al*., 2019). **(B)** Volcano plot on RNA-seq data from CIMP-high vs. non CIMP-high ACC from ACC-TCGA. Light blue dots correspond to differentially expressed genes (adj. p-value=Benjamini-Hochberg-corrected p-value<0.05). Named genes are color-coded by NCI-H295R gene expression percentile (calculated from baseline RNA-seq). *IGF2* is overexpressed in 90% of ACC, not differentially expressed across CIMP classes. **(C)** Total CpG signal across NCI-H295R baseline methylome was summed to predict DNA content, chromosomal segments and copy number, demonstrating “noisy” chromosomal signature characteristic of CIMP-high. In euploidy, Δ Copy number = 0. **(D)** Left, viability curves for NCI-H295R treated with different classes of EZH2i for 96 hours (EPZ-6438, GSK126 are SAM-competitive EZH2i, EED226 is an allosteric EZH2i; n=4 each), x-axis is log10 of the drug concentration in M. Data shown as mean with SEM. Right, western blot measuring H3K27me3 relative to β-actin loading control shown right, n>3; doses tested, GSK126: 0 (Vehicle), 1.25 μM, 5 μM, 7.5 μM, 15 μM, 20 μM (IC-50); EPZ-6438: 0, 1.25 μM, 12.5 μM, 25 μM, 50 μM, 62 μM (IC-50); EED226: 0, 1.25 μM, 12.5 μM, 25 μM, 50 μM, 80 μM (IC-50). **(E)** NCI-H295R transfected with *EZH2* siRNA (siEZH2 A, B, or C) or scrambled negative control (-), and harvested at 144 hours to assess viability (line at mean with 95% CI) and EZH2/H3K27me3/β-actin by western blot (right, n≥2). NCI-H295R doubling time is ∼60 hours; time points selected adequate to measure replication dilution of H3K27me3. **(F)** NCI-H295R were pre-treated with different classes of EZH2i for 96 hours as in D (right), and viable cells were plated at colony forming density in regular (EZH2i-free) medium, grown for 4 weeks. Top, colony area (quantified with Fiji) vs. EZH2i pre-treatment doses. Bottom, representative images of crystal-violet stained colonies at increasing EZH2i doses. Representative experiment shown, n=2. **(G)** *G0S2* methylation after increasing doses of EZH2i (n=3, top) or *EZH2* siRNA (n=1, bottom). Data represented as mean with SEM for each condition. Veh=Vehicle. **(H)** Venn diagram of total number of CpG probes in NCI-H295R methylome and those which were differentially methylated following EZH2i (EPZ-6438 at the IC-50 dose). No probes gained methylation after EZH2i. **(I)** Top, Venn diagram of unique proteins retrieved by DNMT1 IP-MS and EZH2 IP-MS on NCI-H295R nuclear lysates. Bottom, peptides retrieved from DNMT1 or EZH2 IP-MS mapping to known chromatin regulators. SpC=spectral counts. **(J)** Left, heatmap of EZH2, H3K27me3, and H3K27ac ChIP-seq and accessibility (ATAC=ATAC-seq) signal in baseline NCI-H295R at EZH2 peaks, ranked by EZH2 signal. Centered at peak +/-3 kb window. Right, Venn diagram of EZH2, H3K27me3, and H3K27ac peaks. **(K)** Top, Venn diagram of baseline NCI-H295R H3K27me3 peaks and regions targeted for hypermethylation in CIMP-high ACC (DMR+). Bottom, violin plot of average CpG island methylation level (β) in DMR+, DMR+/H3K27me3 overlap regions, and H3K27me3 peaks; lines at median and quartiles. **(L)** Profile plot and heatmap of H3K27me3 signal at regions annotated as baseline NCI-H295R H3K27me3 peaks and DMR+ in NCI-H295R and ENCODE fetal and adult adrenal ChIP-seq (Consortium, 2012; Davis et al., 2018). Centered at peak/DMR +/-3 kb window.

We treated ACC cell lines with S-adenosyl-L-methionine (SAM)-competitive or allosteric EZH2/PRC2 inhibitors (EZH2i). EZH2i induced dose-dependent cell death preceded by H3K27me3 depletion (**Figure 2D, Supp Fig 3H**). In contrast, *EZH2* knockdown induced minimal loss of viability and mild H3K27me3 depletion, suggesting H3K27me3 rather than EZH2 itself is essential for proliferation, **Figure 2E**. Strikingly, transient EZH2i exposure diminished 2D colony formation and survival in a dose-dependent manner (**Figure 2F**), suggesting EZH2i induces heritable epigenomic changes that diminish sustained proliferation potential.

### DNA hypermethylation excludes PRC2 and is associated with aberrant H3K27me3 deposition

We next examined if PRC2 target DNA hypermethylation is directed by catalytically active EZH2 (Viré *et al*., 2006). EZH2i and *EZH2* knockdown did not change DNA methylation at the *G0S2* locus or genome-wide (**Figure 2G-H**). As PRC2 may direct DNMTs through protein-protein interactions (Neri et al., 2013; Viré *et al*., 2006), we performed EZH2-directed complex immunoprecipitation paired with mass spectrometry (IP-MS) and DNMT1 IP-MS on NCI-H295R nuclear lysates. We proceeded with DNMT1 as this is the predominant DNMT in this line, like EZH2 is the predominant H3K27 methyltransferase (**Supp Fig 3I**). DNMT1 also exhibits cell-cycle-dependent upregulation in ACC and cancer (**Supp Fig 1E, 2D**).

We observed EZH2 binds no DNMTs, with virtually no overlap between EZH2 and DNMT1 interactomes (**Figure 2I**). DNMT1 binds many chromatin-bound proteins including the HP1 family of H3K9me readers, representing a conserved DNMT1 mode (Catania et al., 2020), and consistent with a model in which H3K9me3, rather than H3K27me3, instructs CIMP-high DNA methylation (Ohm *et al*., 2007). EZH2 is assembled in PRC2.1, a canonical PRC2 assembly defined by association with PCL accessory proteins that target preferential PRC2 recruitment to unmethylated CpGi (**Figure 2I**, (Li et al., 2017; Perino et al., 2018)).

We next examined EZH2 recruitment genome-wide relative to H3K27me3, active chromatin measured by H3K27 acetylation (H3K27ac), and accessible chromatin. H3K27me3/H3K27ac were mutually exclusive, and most EZH2 peaks co-localized with broad, inaccessible H3K27me3 domains (**Figure 2J**). Regions targeted for hypermethylation in CIMP-high ACC (DMR+) and H3K27me3 peaks exhibited minimal overlap, and DNA methylation levels of H3K27me3 peaks were substantially lower than those of DMR+ (**Figure 2K**). EZH2 and H3K27me3 were excluded from the hypermethylated *G0S2* locus (**Supp Fig 3J**). In fetal and adult adrenals, we observed strong H3K27me3 deposition at DMR+, and reduced deposition at NCI-H295R H3K27me3 peaks (**Figure 2L**). These observations suggest DNA hypermethylation in CIMP-high ACC is propagated independently of PRC2, leads to epigenetic class switching, PRC2 eviction and recruitment to novel sites for H3K27me3 catalysis.

### EZH2i disrupts EZH2 recruitment, wipes H3K27me3, and reverses CIMP-high-defining transcriptional programs

We then treated NCI-H295R cells with EZH2i (EPZ-6438) at the IC-50 dose (**Figure 2D**), and performed chromatin immunoprecipitation sequencing (ChIP-seq). At baseline EZH2 and H3K27me3 sites, EZH2i decreased EZH2 and H3K27me3 while increasing chromatin accessibility. Surprisingly, EZH2i triggered EZH2 displacement to active and accessible chromatin without new H3K27me3 deposition (**Figure 3A**). EZH2i failed to restore expression of hypermethylated genes, like *G0S2*, which was undetectable by RNA-seq at baseline and after EZH2i. However, EZH2i restored expression lowly expressed genes including unmethylated PRC2 targets like *FOXF1* (**Figure 3B; Supp Fig 4A-B**), suggesting catalytically active EZH2 restrains gene expression in CIMP-high ACC. Intriguingly, EZH2i globally disrupted gene expression, with more than half the transcriptome classified as differentially expressed (**Figure 3C**).

**Figure 3:**
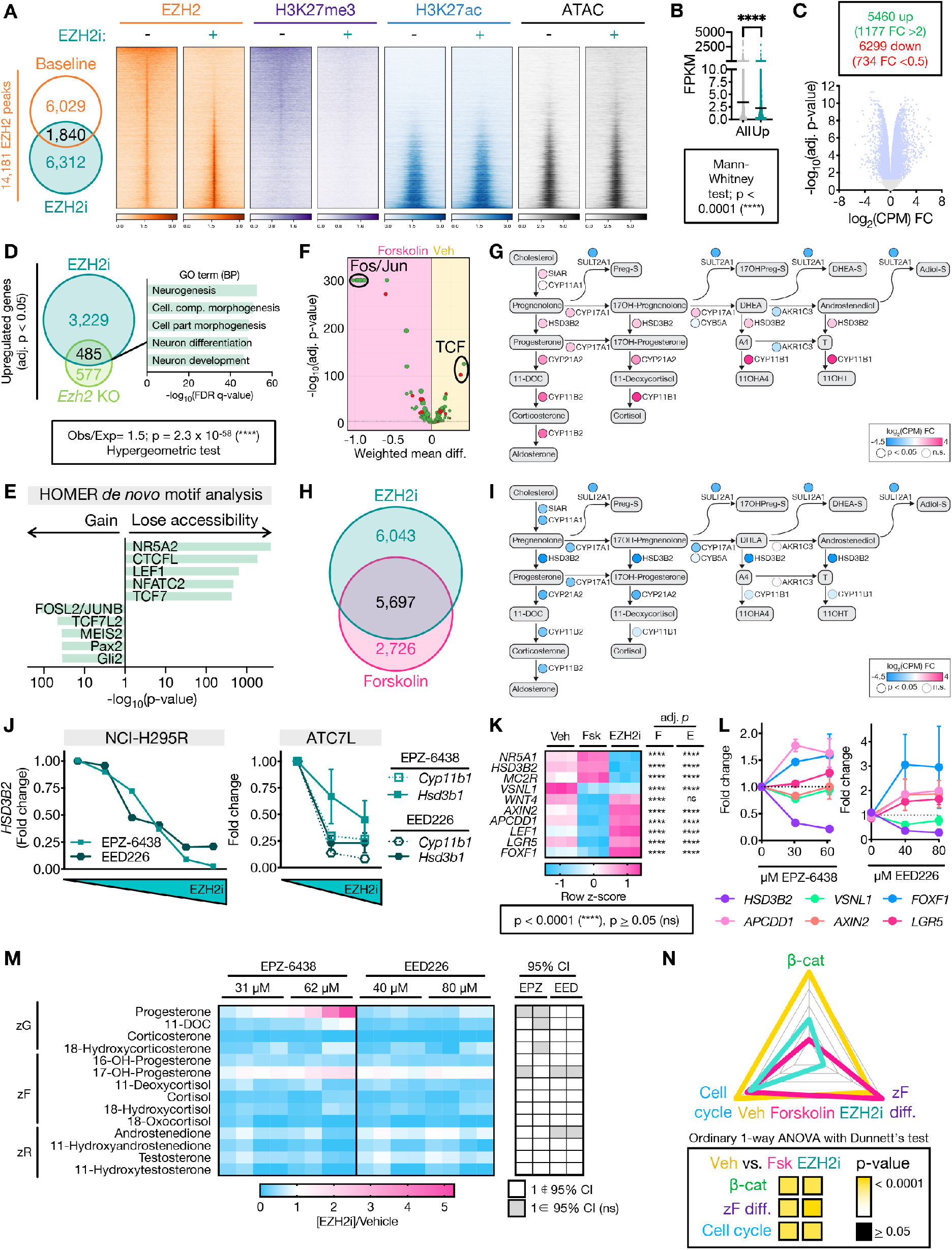
EZH2i disrupts EZH2 recruitment genome-wide, restrains zF differentiation, and reverses the CIMP-high molecular state. **(A)** Left, Venn diagram of NCI-H295R EZH2 peaks at baseline (vehicle-treated, EZH2i-) and after EZH2i (EZH2i+=EPZ-6438 at the IC-50 dose). Right, heatmap of EZH2, H3K27me3, H3K27ac signal in union set of EZH2 peaks at baseline and after EZH2i. Heatmap ranked by ratio of EZH2 signal at baseline to after EZH2i. Centered at peak +/-3 kb window. **(B)** Average baseline NCI-H295R expression (fragments per kilobase of transcript per million mapped reads, FPKM) of all genes in the transcriptome compared to baseline FPKM of genes induced by EZH2i (Up). **(C)** Top, number of differentially expressed genes (adj. p-value<0.05) in EZH2i-vs. vehicle-treated NCI-H295R, FC=fold change. Bottom, corresponding volcano plot. Light blue dots correspond to differentially expressed genes. **(D)** Left, Venn diagram of genes upregulated by EZH2i in NCI-H295R with genes upregulated in mouse model of SF1-driven *Ezh2* deficiency (*Ezh2* KO, (Mathieu *et al*., 2018)). Right, 5 most significant gene sets resulting from GSEA on overlap genes using the GO (BP=Biological Processes) gene set. **(E)** HOMER motif analysis (Heinz et al., 2010) on differentially accessible peaks from NCI-H295R EZH2i compared to baseline ATAC-seq. NR5A2 shares same motif as NR5A1 (SF1). **(F)** DiffTF (Berest et al., 2019) integrating RNA-seq and ATAC-seq from NCI-H295R treated with forskolin to induce zF differentiation vs. vehicle. Negative and positive weighted mean difference reflect transcription factor signal stronger in forskolin- or vehicle-treated cells, respectively. **(G)** Steroidogenesis diagram depicting impact of forskolin on expression of zonally expressed steroidogenic enzymes in NCI-H295R by RNA-seq. **(H)** Venn diagram of differentially expressed genes (compared to baseline) in NCI-H295R treated with EZH2i or forskolin by RNA-seq. **(I)** Steroidogenesis diagram depicting impact of EZH2i on enzyme expression in NCI-H295R by RNA-seq. **(J)** Fold change in expression of steroidogenic enzymes in ACC cell lines treated with increasing doses of EZH2i measured by qPCR. NCI-H295R, n=1. ATC7L, n=2-3 for all points. Data shown as mean with SEM. Concentrations tested were: NCI-H295R, as in **Figure 1D**; ATC7L - 0, 42 μM, 84 μM EPZ-6438 (IC-50) and 0, 53 μM, 107 μM EED226 (IC-50). **(K)** Heatmap of gene expression of NCI-H295R at baseline (vehicle, Veh), following forskolin (Fsk) treatment or following EZH2i with adj. p-value for comparison to Veh shown right, by RNA-seq. *FOXF1* is a PRC2 target (**Supp Fig 4B**). Per **Supp Fig 3B** and (Nishimoto et al., 2015), “zF genes”= *HSD3B2, MC2R* and remainder are “zG genes”. **(L)** NCI-H295R were treated with indicated doses of EZH2i followed by forskolin. Gene expression measured by qPCR, n=3. Data shown as mean with SEM. **(M)** LC-MS/MS analysis of media from NCI-H295R treated as in L to measure zone-specific steroid output. Left, heatmap depicting normalized steroid output relative to vehicle. Right, significance of change. If the null value (1) falls in the 95% CI of the mean for each treatment group, change in steroid output is considered insignificant (ns). **(N)** ZF differentiation, Wnt, and cell cycle scores for NCI-H295R at baseline (Veh) or treated with forskolin (Fsk) or EZH2i, calculated and graphed as in **Figure 1B**.

ACC exhibit a spectrum of zF differentiation, Wnt/β-catenin activation and proliferation, with CIMP-high ACC at the relative maxima of these poles (**Figure 1B; Supp Fig 1, 3B-G**). In the mouse model of SF1-driven *Ezh2* ablation, mice develop glucocorticoid insufficiency due to a failure of the zF to differentiate and proliferate in response to ACTH (Mathieu et al., 2018). Despite H3K27me3 deposition at several new sites in CIMP-high ACC (**Figure 2L**), EZH2i derepressed >50% of the genes induced in this mouse model including neuronal programs (**Figure 3D**). This was accompanied by marked epigenetic changes, where EZH2i (1) restored chromatin accessibility of programs silenced in steroidogenic adrenocortical cells (e.g. those driven by PAX2 and GLI), (2) disrupted accessibility of targets of canonical Wnt/β-catenin transcription factors TCF/LEF, and (3) reduced accessibility of putative SF1 and CTCFL targets (**Figure 3E**). Based on these findings, we hypothesized EZH2i disrupts adrenocortical differentiation.

To evaluate this, we treated NCI-H295R cells with forskolin, a zF differentiation agent that induces the PKA/cAMP signaling cascade downstream of ACTH (Seamon et al., 1981; Xing et al., 2011). Forskolin administration (1) increased expression of zF differentiation genes, (2) diminished expression and accessibility of TCF targets, and (3) induced expression of zonally expressed steroidogenic enzymes (**Figure 3F-G; Supp Fig 4C**). Strikingly, EZH2i disrupted ∼70% of genes differentially expressed after forskolin treatment (**Figure 3H**), and potently downregulated steroidogenic enzymes (**Figure 3I**). Suppression of steroidogenic enzymes was dose-dependent and observed with different classes of EZH2i across ACC cell lines (**Figure 3J**). Moreover, EZH2i pretreatment followed by forskolin administration diminished both forskolin-induced silencing of canonical Wnt targets and induction of steroidogenic enzymes, ultimately restraining steroid output (**Figure 3K-M**).

These observations are consistent with a role for EZH2 in programming the cellular response to ACTH/PKA in CIMP-high ACC, though not solely by dysregulating expression of PKA signaling components (Mathieu *et al*., 2018). Though EZH2i induced few canonical Wnt targets (**Figure 3E, K**), EZH2i reversed all three core modules of CIMP-high ACC (**Figure 3N**), while forskolin induced differentiation at the expense of cell cycle and Wnt activation (**Figure 3F, K, N**). Observing that EZH2i disrupted a spectrum of transcriptional programs (**Figure 3C**), including those governed by EZH2 in the normal adrenal (**Figure 3D**), led us to explore if the effects of EZH2i result from an alternative EZH2 role.

### EZH2 binds transcriptional coactivator β-catenin in an off-chromatin complex

EZH2 IP-MS revealed several non-PRC2 partners, including nuclear receptors known to regulate adrenocortical biology (Bassett et al., 2004), as well as β-catenin (**Figure 4A**), constitutively active in NCI-H295R due to the p.S45P mutation (**Supp Fig 3A**). Given β-catenin’s abundance in the EZH2 interactome (**Figure 4A**), possible role in the chromatin response to EZH2i (**Figure 3E**) and its well-established role in adrenocortical differentiation and tumorigenesis, we elected to focus our studies on EZH2/β-catenin.

**Figure 4:**
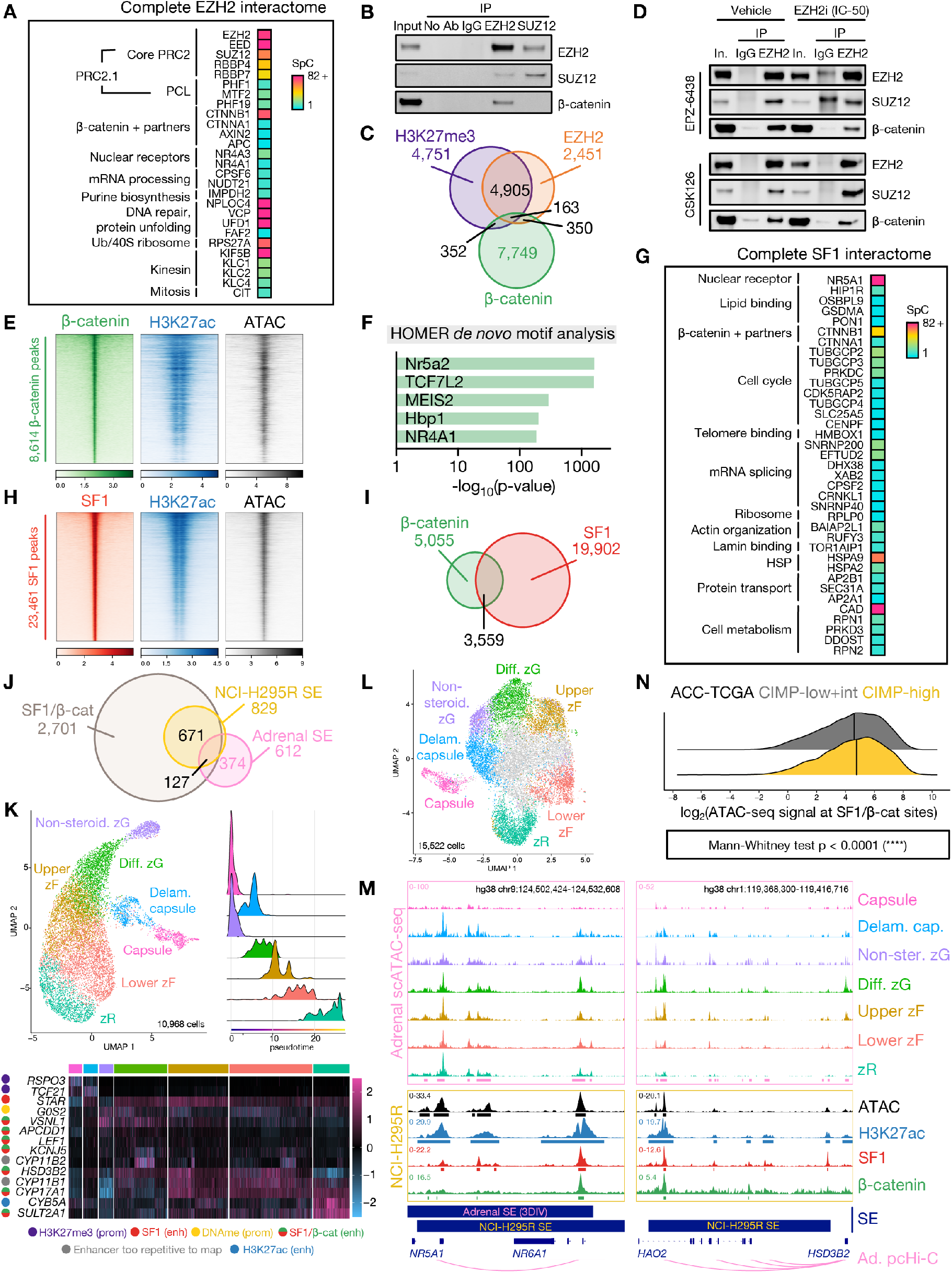
Nuclear pools of off-chromatin (EZH2-bound) and on chromatin (SF1-bound) β-catenin direct upper zF differentiation in CIMP-high ACC. **(A)** Peptides retrieved from EZH2 IP-MS on NCI-H295R nuclear lysates. **(B)** Representative western blot of NCI-H295R nuclear co-IP, detecting EZH2, SUZ12 and β-catenin. Lanes are 10% input, negative control co-IP (no antibody, IgG), EZH2 IP and SUZ12 IP. n>5 (EZH2 IP), n=2 (SUZ12 IP). **(C)** Venn diagram of H3K27me3, EZH2, and β-catenin ChIP-seq peaks in baseline NCI-H295R. **(D)** Representative western blot of EZH2 IP in vehicle-(left) or EZH2i-treated (right, IC-50 dose) NCI-H295R. Lanes are 10% input, negative control IgG IP, EZH2 IP. Higher molecular weight band in EPZ-6438 IgG lane in EZH2/SUZ12 blots is non-specific and emerges when using the same antibody species for IP and western. **(E)** Heatmap of β-catenin, H3K27ac, and ATAC signal in baseline NCI-H295R at β-catenin peaks, ranked by β-catenin signal. Centered at peak +/-3 kb window. **(F)** HOMER motif analysis on baseline NCI-H295R β-catenin peaks. **(G)** Peptides retrieved from SF1 IP-MS on NCI-H295R nuclear lysates. **(H)** Heatmap of SF1, H3K27ac and ATAC signal in baseline NCI-H295R at SF1 peaks, ranked by SF1 signal. Centered at peak +/-3 kb window. **(I)** Venn diagram of baseline NCI-H295R SF1 and β-catenin peaks. **(J)** Venn diagram of baseline NCI-H295R SF1/β-catenin peaks, baseline NCI-H295R super-enhancers (SE), and physiologic adrenal SE called by 3DIV (Kim et al., 2020; Yang et al., 2018) on ENCODE samples. **(K)** Single-cell RNA-seq (scRNA-seq) data from fetal, neonatal and adult human adrenal (Han *et al*., 2020) was integrated and analyzed with comparison to reference markers to identify populations comprising the corticocapsular unit. Top left, scRNA-seq UMAP. Non-steroid.=Non-steroidogenic, Diff.=Differentiated, Delam.=Delaminating. Top right, scRNA-seq pseudotime analysis (origins set for fetal and adult adrenal populations in the capsule and non-steroidogenic zG). Bottom, heatmap of scaled expression of lineage-defining genes across scRNA-seq with epigenetic regulation in NCI-H295R shown left. Prom=promoter, enh=enhancer, DNAme=DNA methylation. Gene regulation by active (H3K27ac only) and SF1- or SF1/β-catenin-bound enhancers was identified by overlap of ChIP-seq with adrenal promoter capture Hi-C (pcHi-C (Jung *et al*., 2019)). **(L)** Single-cell ATAC-seq (scATAC-seq) data from fetal and adult human adrenal (Domcke *et al*., 2020; Zhang *et al*., 2021) was integrated and analyzed with comparison to scRNA-seq and reference markers to identify analogous populations comprising the corticocapsular unit. Cells colored in grey in UMAP plot are likely cortical given accessibility within *NR5A1* locus though possess ambiguous classification. **(M)** Example adrenal scATAC-seq and NCI-H295R ATAC, SF1, β-catenin, and H3K27ac tracks across the *NR5A1* and *HSD3B2* loci in baseline NCI-H295R. Adrenal scATAC peak calls are depicted by pink bars at the bottom of top window, and NCI-H295R peak calls are depicted by bars below each track. 3DIV annotation of adrenal super-enhancers (SE), and NCI-H295R baseline SE are shown by bars below window. Promoter/enhancer contacts from adrenal (Ad.) pcHi-C (Jung *et al*., 2019) are depicted below gene annotations. **(N)** Ridge plot of chromatin accessibility signal at SF1/β-catenin co-targets in ACC-TCGA ATAC-seq samples (n=9, (Corces *et al*., 2018)). Line at median.

We were unable to purify endogenous β-catenin-containing transcriptional complexes by β-catenin IP-MS, due to β-catenin’s strong affinity for contaminating adherens junctions (Yakulov et al., 2013), **Supp Fig 4D**. However, EZH2, SUZ12, and β-catenin co-eluted with histones bearing K27 modifications by size exclusion chromatography (SEC) (**Supp Fig 4E**). While EZH2 IP consistently retrieved both SUZ12 (a core PRC2 member) and β-catenin, SUZ12 IP did not retrieve β-catenin (**Figure 4B**). EZH2/H3K27me3/β-catenin possess minimal overlap on chromatin (**Figure 4C**), and EZH2i did not disrupt nuclear EZH2/β-catenin (**Figure 4D**). Together, these data suggest EZH2/β-catenin is a nuclear but off-chromatin complex that may compete for β-catenin binding to chromatin and participate in the epigenetic response to EZH2i, in contrast to its role in other contexts (Hoffmeyer et al., 2017).

EZH2i reversed the transcriptional and epigenomic programs that define CIMP-high ACC (**Figure 3E, 3N**). Given CIMP-high are Wnt-active and β-catenin is a major EZH2 binding partner spared by EZH2i, we next investigated β-catenin’s role on chromatin.

### SF1/β-catenin regulates a super-enhancer-driven zF differentiation program in ACC

In NCI-H295R cells, β-catenin colocalized with active and accessible chromatin regions genome-wide (**Figure 4E**). Surprisingly, β-catenin peaks exhibited substantial enrichment for the SF1 motif, comparable to enrichment for TCF/LEF motifs (**Figure 4F**). This was consistent with SEC demonstrating co-elution of SF1, β-catenin, and H3K27ac in ACC cell lines (**Supp Fig 4E**). By SF1 IP-MS, we observed that p.S45P β-catenin is a major SF1 binding partner in NCI-H295R cells (**Figure 4G**).

We and others previously reported an SF1/β-catenin complex, thought to regulate gene expression but without a clear global role (Gummow et al., 2003; Hossain and Saunders, 2003; Kennell et al., 2003; Mizusaki et al., 2003). ChIP-seq for SF1 in NCI-H295R cells revealed that, as expected, SF1 binds active and accessible chromatin regions (**Figure 4H**). There was substantial overlap between SF1 and β-catenin recruitment (**Figure 4I**), and SF1/β-catenin sites encompassed predominantly distal regions (**Supp Fig 4F**), suggesting enhancer regulation.

A special class of enhancers, super-enhancers (SE), are defined by high density H3K27ac and possess occupancy by lineage-defining transcription factors, driving cell-of-origin programs in development and disease (Hnisz et al., 2013; Hnisz et al., 2015; Lovén et al., 2013; Pott and Lieb, 2015; Sabari et al., 2018; Whyte et al., 2013; Zamudio et al., 2019). We observed that >90% of normal adrenal SE remain active enhancers in NCI-H295R cells (**Supp Fig 4G**); however, the vast majority of NCI-H295R SE are novel (**Figure 4J**). Strikingly, virtually all NCI-H295R SE possess SF1/β-catenin, representing a major departure from normal adrenal (**Figure 4J**).

To understand the genomic architecture and physiologic relevance of these SE, we examined promoter capture Hi-C, single-cell RNA-seq, and single-cell ATAC-seq from normal adrenals (Domcke et al., 2020; Han et al., 2020; Jung et al., 2019; Zhang et al., 2021), and integrated these studies with our epigenomics data. We observed SF1/β-catenin SE and enhancers contact promoters of many genes critical for zone-specific and adrenocortical steroidogenic identity, including *HSD3B2* and *NR5A1* itself (**Figure 4K-M**). These enhancers are present in normal adrenal (**Figure 4M, Supp Fig 4H**), and more prominent in outer cortex, consistent with nuclear localization of β-catenin in these regions (**Figure 1A, 4L-M**). Genes that retain promoter H3K27me3 in NCI-H295R cells (e.g. capsular genes *RSPO3* and *TCF21*) are also silenced in the cortex (**Figure 4K**).

To extend these observations to ACC, we analyzed ACC-TCGA ATAC-seq (Corces et al., 2018) and observed CIMP-high have increased accessibility of SF1/β-catenin co-targets (**Figure 4N**). Collectively, these data suggest epigenetic programming in CIMP-high ACC maintains a differentiation state resembling the upper zF through SF1/β-catenin chromatin regulation that co-opts physiologic programming.

### EZH2/β-catenin and SF1/β-catenin are selected for through all stages of adrenocortical neoplasia

Members of our team recently developed an autochthonous mouse model of zF-differentiated ACC, driven by combined β-catenin gain-of-function (GOF)/p53 loss-of-function (LOF) in cells expressing adrenocortical enzyme Cyp11b2, BPCre^AS/+^ (Borges et al., 2020). β-catenin GOF causes adrenal hyperplasia, though single hit β-catenin GOF or p53 LOF is insufficient for ACC (Borges *et al*., 2020; Pignatti et al., 2020). Genetically, BPCre^AS/+^ is similar to CIMP-high ACC, and exhibits selective expansion, unrestricted growth, and dissemination of β-catenin/cell cycle-avid cells with high EZH2 expression and autonomous glucocorticoid production (**Supp Fig 5A, Figure 5A-B**, (Borges *et al*., 2020)).

**Figure 5:**
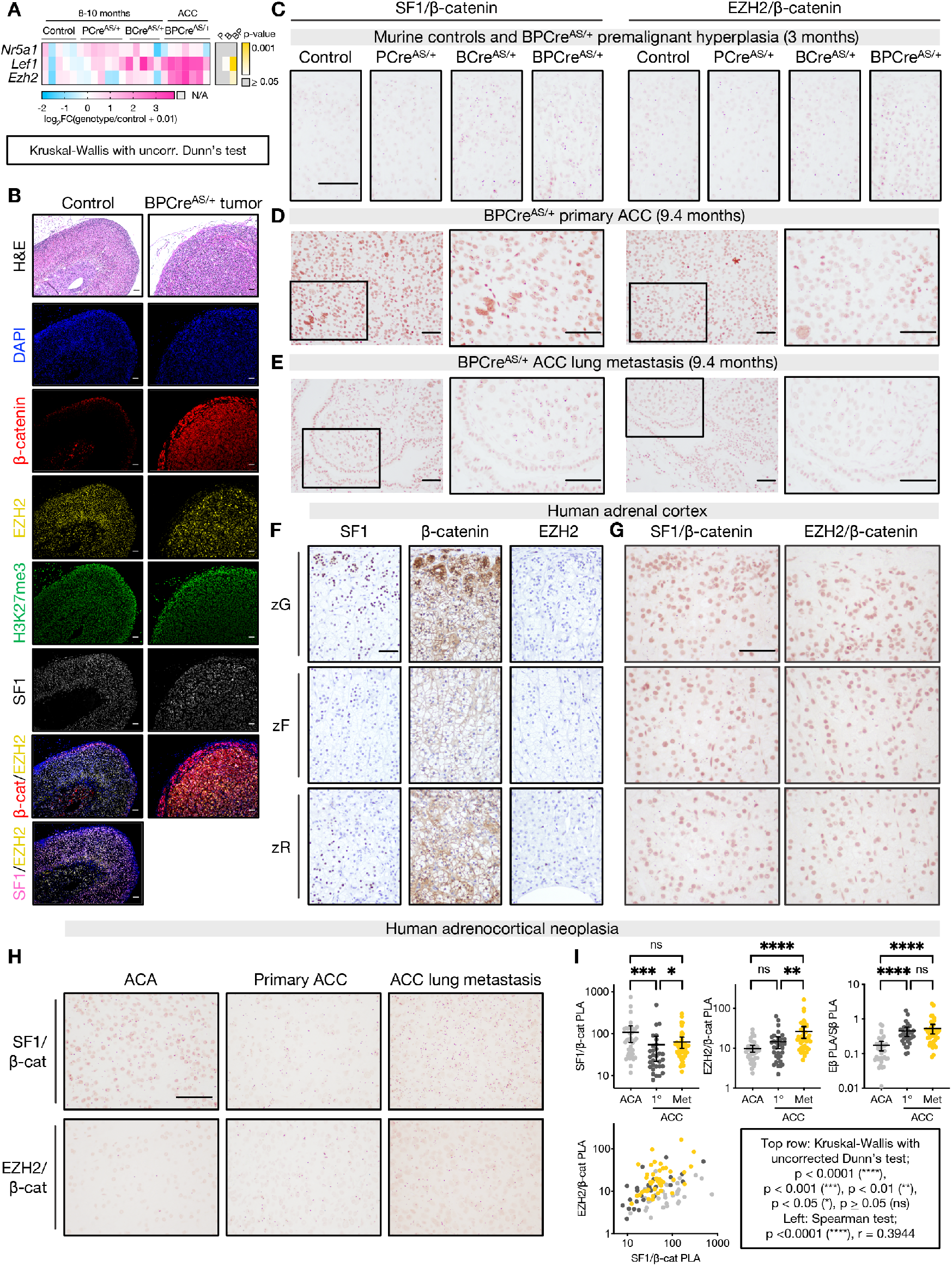
Nuclear EZH2/β-catenin and SF1/β-catenin complexes persist through adrenocortical neoplastic evolution. **(A)** Left, heatmap of gene expression measured by qPCR in adrenals from control mice (AS^Cre/+^), mice with p53 LOF (PCre^AS/+^), mice with β-catenin GOF (BCre^AS/+^), or ACC from combined p53 LOF/β-catenin GOF (BPCre^AS/+^). Right, p-value for each genotype compared to control. **(B)** Top row, representative hematoxylin and eosin (H&E) staining of control adult mouse adrenal and BPCre^AS/+^ primary tumor (10-month-old). Rows 2-6, immunofluorescence staining nuclei (DAPI), β-catenin, EZH2, H3K27me3, or SF1. Rows 7-8, colocalization of β-catenin or SF1 and EZH2. Bar=50 μm. **(C-E)** Representative images of SF1/β-catenin (left) and EZH2/β-catenin (right) proximity ligation assay (PLA) performed on 3-month-old adrenals (control, n=4; PCre^AS/+^, n=3; BCre^AS/+^, n=3; BPCre^AS/+^, n=4), BPCre^AS/+^ primary ACC (n=5), BPCre^AS/+^ lung metastases (n=3). PLA signal are subnuclear pink dots. Bar=100 μm. **(F-G)** Representative images of SF1, β-catenin, and EZH2 IHC or SF1/β-catenin and EZH2/β-catenin PLA in human adult adrenal cortex (n=2). Bar=100 μm. **(H)** Representative images of SF1/β-catenin and EZH2/β-catenin PLA in a TMA of human adult ACA (n=39) and primary (n=34) and metastatic (n=35) ACC. **(I)** Normalized TMA PLA signal, quantified by Fiji. Each sample is represented by a point. Top right, “Eβ PLA/Sβ PLA” refers to ratio of EZH2/β-catenin PLA signal to SF1/β-catenin PLA signal. Top row, line at mean with 95% CI.

To determine if EZH2/β-catenin and SF1/β-catenin complexes participate in tumorigenesis, we optimized a technique to detect protein-protein interactions *in situ* (proximity ligation assay, PLA). We observed that β-catenin-containing complexes are nuclear, and zonally distributed in control mice (**Figure 5C, Supp Fig 5B-C**), mirroring the Wnt/β-catenin signaling gradient (**Figure 1A**). We also observed transformation and metastatic seeding of cells uniformly expressing EZH2/β-catenin and SF1/β-catenin complexes (**Figure 5D-E, Supp Fig 5D**).

Given the persistence of EZH2/β-catenin and SF1/β-catenin complexes throughout murine carcinogenesis and the increased accessibility of SF1/β-catenin co-targets in CIMP-high ACC (**Figure 4N**), we speculated that these complexes would accompany human carcinogenesis. We applied PLA to the normal human adrenal cortex, benign adrenocortical tumors, and primary and metastatic ACC. In the human adrenal, we observed abundant SF1/β-catenin complexes following the Wnt/β-catenin gradient, and infrequent EZH2/β-catenin complexes reflecting the rarity of EZH2 expression (**Figure 5F-G**). In human adrenocortical tumors, we observed retention of β-catenin-containing complexes through metastatic disease (**Figure 5H-I**), with increased abundance of EZH2/β-catenin complexes relative to SF1/β-catenin complexes in malignancy (**Figure 5I**) mirroring increased EZH2 expression in cancer (**Figure 1G**). These data demonstrate that persistence of EZH2/β-catenin and SF1/β-catenin complexes is conserved across murine and human adrenocortical carcinogenesis, suggesting programs correlated with or coordinated by these complexes are subject to positive selection through all phases of CIMP-high ACC evolution.

### EZH2i erases SF1/β-catenin-dependent transcriptional and epigenetic programming

The presence of EZH2/β-catenin and SF1/β-catenin *in vivo* was compelling, given EZH2/β-catenin is an off-chromatin complex that persists with EZH2i (**Figure 4B-D**). We also identified that EZH2i reverses zF differentiation (**Figure 3**), coordinated by SF1/β-catenin in CIMP-high ACC (**Figure 4E-N**). Furthermore, nearly 40% of genes putatively regulated by SF1/β-catenin enhancers are downregulated by EZH2i (**Figure 6A**).

**Figure 6:**
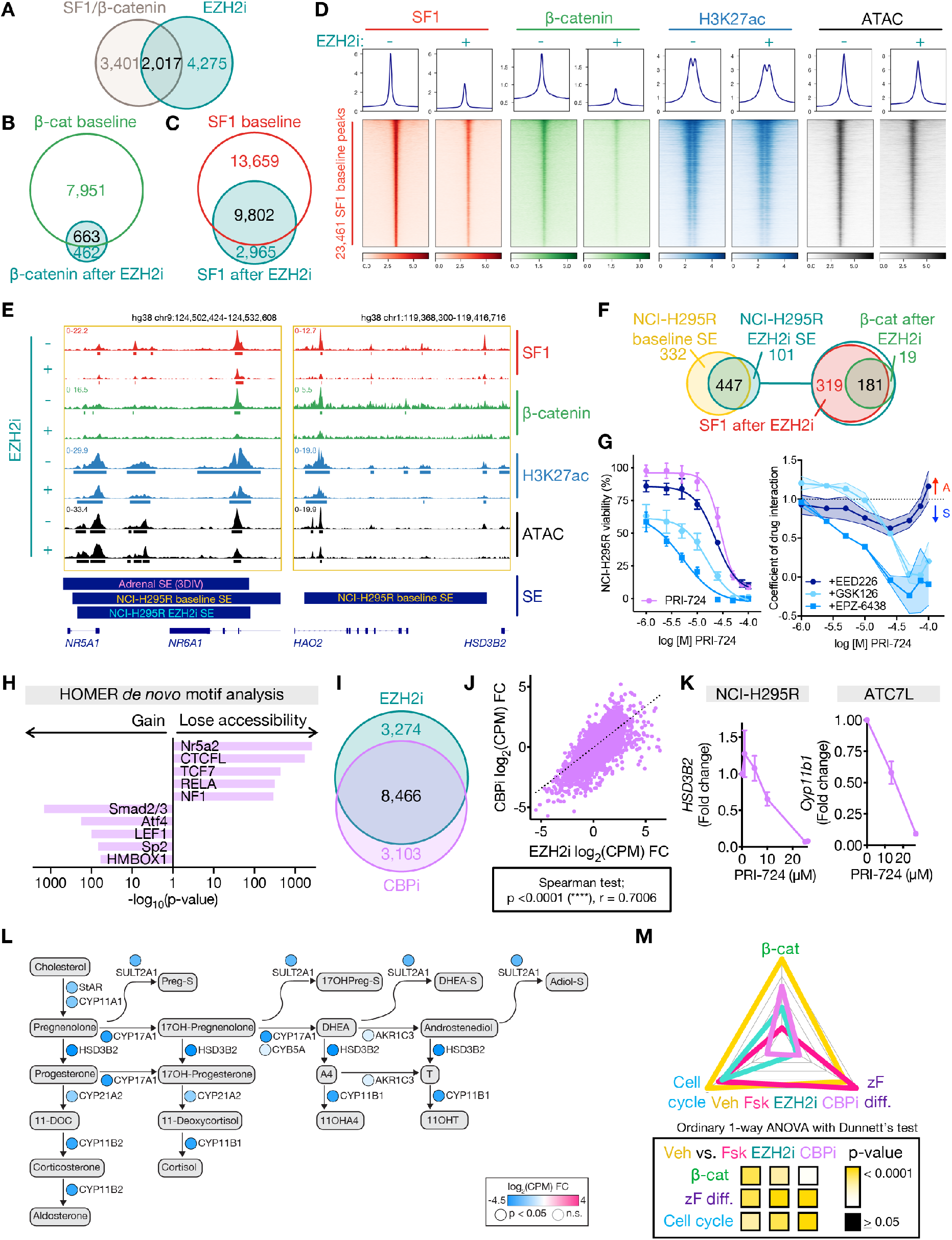
EZH2i evicts SF1 and β-catenin genome-wide, disrupting enhancer programming in CIMP-high ACC. **(A)** Venn diagram of genes putatively regulated by active SF1/β-catenin-bound enhancers and downregulated by EZH2i. **(B-C)** Venn diagram of NCI-H295R β-catenin or SF1 peaks at baseline (vehicle-treated) and after EZH2i. **(D)** Profile plot and heatmap of NCI-H295R SF1, β-catenin, H3K27ac, and ATAC signal at SF1 peaks at baseline (EZH2i-) and after EZH2i (EZH2i+), ranked by baseline SF1 signal. Centered at peak +/-3 kb window. **(E)** Example SF1, β-catenin, H3K27ac ChIP-seq, and accessibility (ATAC-seq) tracks across the *NR5A1* and *HSD3B2* loci in NCI-H295R at baseline (as in **Figure 4M**) or after EZH2i. Peak calls depicted by bars below each track. 3DIV annotation of adrenal SE, NCI-H295R baseline and EZH2i SE shown by bars below window. **(F)** Left, Venn diagram of NCI-H295R SE at baseline and after EZH2i. Right, Venn diagram of NCI-H295R EZH2i SE with SF1 EZH2i peaks and β-catenin EZH2i peaks. **(G)** Left, viability curves for NCI-H295R treated with increasing concentrations of CBP inhibitor (CBPi) PRI-724 +/-different EZH2i (GSK126, EPZ-6438 or EED226) at the IC-50 dose. Right, coefficient of drug interaction (CDI) for viability. CDI>1 represents antagonism (A), CDI <1 represents synergy (S). Data are represented by mean with SEM (whiskers or error bands). CBPi, n=9; CBPi + GSK126, n=3; CBPi + EPZ-6438, n=3; CBPi + EED226, n=3. **(H)** HOMER motif analysis on differentially accessible peaks from NCI-H295R CBPi (IC-50) compared to baseline ATAC-seq. **(I)** Venn diagram of differentially expressed genes (compared to baseline) in NCI-H295R treated with EZH2i or CBPi (IC-50) measured by RNA-seq. **(J)** Scatterplot of change in gene expression (compared to baseline) in NCI-H295R treated with CBPi vs. EZH2i. **(K)** Fold change in expression of steroidogenic enzymes in ACC cell lines treated with increasing doses of CBPi measured by qPCR. NCI-H295R, n=2. ATC7L, n=3. Data shown as mean with SEM. **(L)** Steroidogenesis diagram depicting impact of CBPi on enzyme expression in NCI-H295R by RNA-seq. **(M)** ZF differentiation, Wnt, and cell cycle scores for NCI-H295R at baseline (Veh) or treated with forskolin (Fsk), EZH2i, or CBPi calculated and graphed as in **Figure 3N**.

Strikingly, EZH2i evicted SF1 and β-catenin genome-wide, decreasing H3K27ac and accessibility at SF1 sites (**Figure 6B-D**). We speculated EZH2i expunges β-catenin from chromatin via accumulation of EZH2/β-catenin (**Figure 4D**) induced by EZH2 eviction from H3K27me3 domains (**Figure 3A**). As EZH2 and SF1 do not directly interact (**Figures 4A, 4G**), it was difficult to rationalize why EZH2i disrupted global SF1 programming even at regions not annotated as β-catenin co-targets (**Figure 6C-D**). However, EZH2i disrupted SF1/β-catenin recruitment to prototype SE including those regulating *NR5A1* (encoding SF1) and *HSD3B2*, resulting in decreased gene expression and accessibility (**Figure 3J-K, 6E**). Indeed, EZH2i downgraded half of all baseline SE (**Figure 6F**).

To assess the extent to which EZH2i genomic changes result directly from SE disruption, we treated ACC cell lines with a specific, irreversible inhibitor of CBP (CBPi PRI-724 (Kahn, 2014)), an H3K27 acetyltransferase required for enhancer activity (Merika et al., 1998; Raisner et al., 2018). CBPi is unlikely to directly alter EZH2 recruitment; however, CBPi induced dose-dependent loss of viability synergistic with EZH2i in all ACC cell lines (**Figure 6G; Supp Fig 5E-F**). We observed redundant and highly correlated effects of CBPi and EZH2i on the NCI-H295R epigenome (**Figure 6H**) and transcriptome (**Figure 6I-J**), including dose-dependent dedifferentiation (**Figure 6K-L**). Like EZH2i, CBPi reverses all core modules that define CIMP-high ACC (**Figure 6M**). Together, these data suggest SF1/β-catenin enhancer programming is a central CIMP-high susceptibility.

### EZH2i hinders tumor growth, proliferation, and SF1/β-catenin-dependent differentiation *in vivo*

To determine if differentiation programs targeted by EZH2i represent a viable therapeutic strategy in zF-differentiated ACC bearing SF1/β-catenin and EZH2/β-catenin complexes, we developed a subcutaneous allograft model using a cell line (BCH-ACC3A) derived from BPCre^AS/+^ ACC with metastatic potential, **Figure 7A**. Recipient mice were treated with vehicle or EZH2i, which was well tolerated (**Supp Fig 5G**). EZH2i-treated mice exhibited diminished tumor growth (**Figure 7B**) with decreased H3K27me3 deposition (**Figure 7C**) and proliferation (**Figure 7D, Supp Fig 5H**). Tumors from EZH2i-treated mice also exhibited dedifferentiation (**Figure 7E**) with a reduction in SF1/β-catenin complexes (**Figure 7F**) and retention of EZH2/β-catenin complexes (**Figure 7G**), recapitulating the molecular features we observed *in vitro* (**Figures 3-4, 6**). Taken together, these studies point to zF differentiation as a targetable epigenetic vulnerability selected for in CIMP-high ACC and provide proof-of-principle support for efficacy of dedifferentiating therapies in ACC treatment.

**Figure 7:**
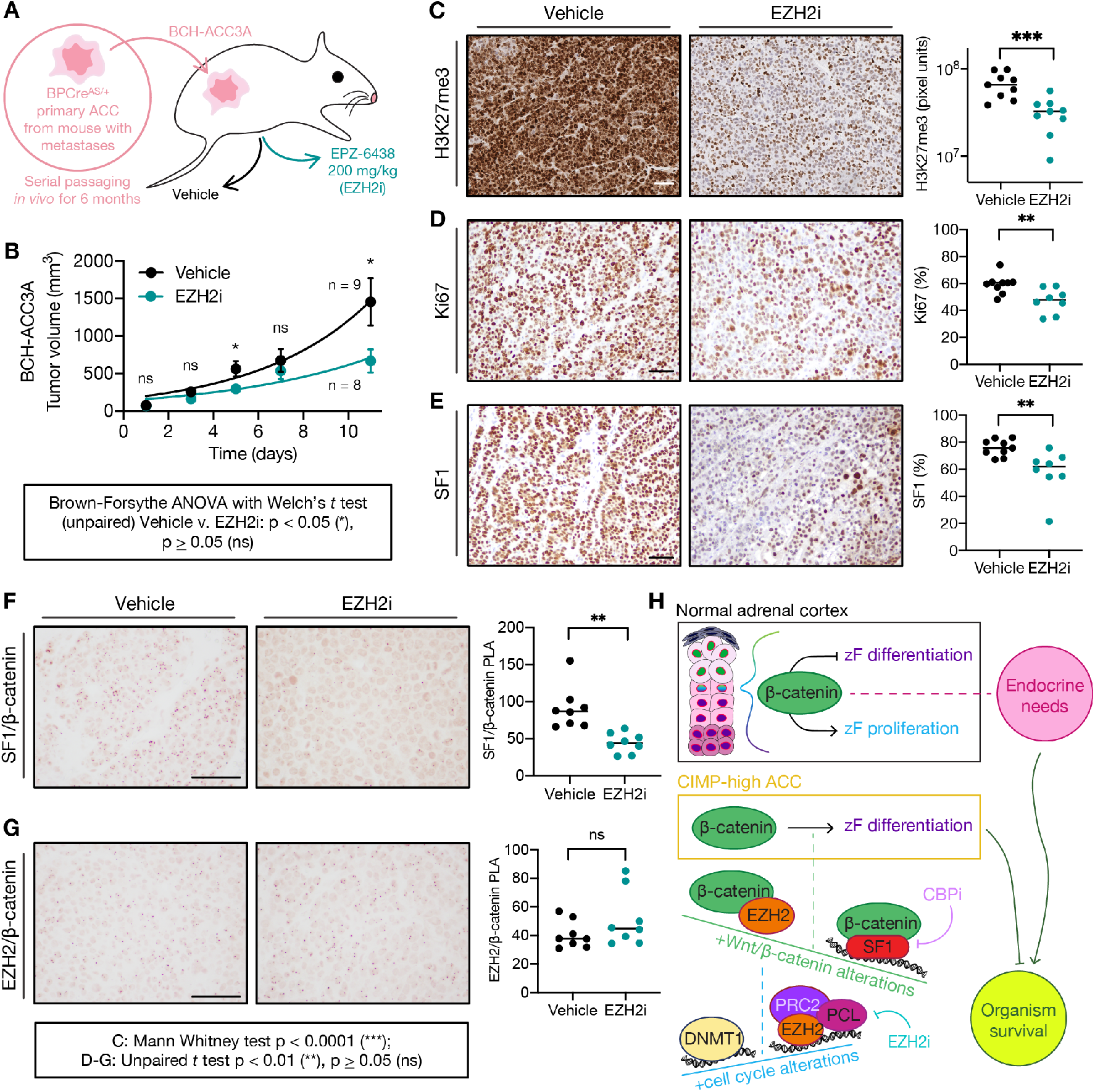
EZH2i hinders ACC growth, proliferation, and differentiation *in vivo*. **(A)** Derivation of BCH-ACC3A cell line and subcutaneous NSG mouse allograft model, randomized to vehicle or EZH2i treatment at tumor volume 100 mm^3^. **(B)** Tumor growth across treatment groups; data shown as mean with SEM. **(C)** H3K27me3 IHC across treatment groups. Left, bar=50 μm; right, each sample represented by 3 points, H3K7me3 quantified by MATLAB, line at median. **(D-G)** Ki67 IHC, SF1 IHC, SF1/β-catenin PLA or EZH2/β-catenin PLA across treatment groups. Left, bar=100 μm; right, each sample is represented by a point and % nuclear signal or normalized PLA signal quantified by Fiji, line at median. **(H)** Model: In the upper zF of the normal adrenal cortex, β-catenin restrains zF differentiation or permits zF proliferation depending on endocrine demands (systemic need for glucocorticoids and flux through ACTH). This homeostasis is required for organism survival. In CIMP-high ACC, β-catenin drives zF differentiation through SF1/β-catenin hijacking of genome-wide SE. SF1/β-catenin’s actions on chromatin are limited by EZH2/β-catenin, an off-chromatin complex that completes for β-catenin binding. EZH2/β-catenin abundance is limited by on chromatin EZH2 and PRC2 catalytic activity. PRC2 remains catalytically active in CIMP-high ACC despite displacement by CpGi hyper-methylation (written by DNA methyltransferases like DNMT1). Recurrent Wnt pathway and cell cycle alterations in CIMP-high ACC promote the formation of β-catenin-containing and EZH2-containing complexes. Ultimately, β-catenin-dependent zF differentiation is required for sustained ACC proliferation at the cost of organism survival. This program is erased by ACC dedifferentiating agents like EZH2i or CBPi, representing a promising therapeutic avenue.

## DISCUSSION

ACC is exquisitely rare and outcomes remain dismal. CIMP-high ACC is prevalent, invariably metastatic, and lethal (Mohan *et al*., 2019; Zheng *et al*., 2016). Molecularly, CIMP-high ACC is defined by abnormal DNA methylation and paradoxical activation of cell cycle, Wnt/β-catenin signaling, and zF differentiation. We discovered that abnormal CpGi hypermethylation, in addition to serving as a pathognomonic marker of aggressive disease, displaces EZH2/PRC2 to novel sites. CIMP-high DNA hypermethylation is uniform, with many targets possessing binary and complete methylation (Mohan *et al*., 2019). These data, consistent with literature examining etiology and emergence of CIMP-high (Tao *et al*., 2019; Vaz *et al*., 2017), suggest acquisition of this signature is an early selection event in adrenocortical carcinogenesis.

We reconcile the convergence of zF differentiation and Wnt/β-catenin activation in ACC by discovering uniform SF1/β-catenin control of CIMP-high SE, including an SE that regulates expression of SF1 itself. Selection for zF differentiation is counterintuitive given the many events other cancers acquire to harness proliferation potential at the expense of differentiation (Ireland et al., 2020; LaFave et al., 2020). Our data suggest that the cell originating CIMP-high ACC is one which relies on β-catenin and zF differentiation for sustained proliferation, perhaps a transit-amplifying ACTH-responsive cell of the upper zF which fails to proliferate in the setting of congenital adrenocortical EZH2 deficiency or β-catenin ablation (Finco et al., 2018; Mathieu *et al*., 2018). This is further supported by the demonstrations that combined β-catenin/cell cycle GOF generates zF-differentiated glucocorticoid producing ACC (Borges *et al*., 2020), and adrenocortical deficiency of negative Wnt pathway regulator ZNRF3 induces zF hyperplasia with eventual malignant transformation (Basham et al., 2019; Warde et al., 2022; Wilmouth et al., 2022).

We identify a series of protein complexes, EZH2/β-catenin and SF1/β-catenin, that shuttle β-catenin off and on chromatin. These complexes are zonally distributed, conserved across murine and human adrenocortical carcinogenesis, and selected for throughout ACC evolution. EZH2/β-catenin formation is a necessary but costly consequence of Wnt and cell cycle activation in CIMP-high, destabilizing the SF1/β-catenin program. CIMP-high ACC may thus require high PRC2 catalytic activity to restrain EZH2/β-catenin in the setting of CpGi hypermethylation, in contrast to other tissues that use CIMP-high to select for PRC2 loss of function with malignancy (Bayliss et al., 2016). The EZH2/β-catenin/SF1 triangulation therefore creates a dependence of zF differentiation on PRC2, illuminating an intrinsic tissue-specific vulnerability in CIMP-high ACC with therapeutic significance *in vitro* and *in vivo* (**Figure 7H**).

Repressive epigenetic modifiers are often modeled as complexes that maintain stemness. PRC2 is critical for embryonic pluripotency and gastrulation (Deevy and Bracken, 2019), and in many cancers restrains differentiation for sustained proliferation potential (Schuettengruber *et al*., 2017), in apparent contrast to our work. A perhaps more nuanced interpretation is that PRC2/H3K27me3 deposition facilitates cell state transitions required for accurate differentiation. Indeed, our studies reveal that catalytically active PRC2 stabilizes a pro-proliferative differentiation state in CIMP-high ACC, by limiting EZH2’s interaction with a transcriptional coactivator core to adrenocortical cell type specification, β-catenin.

The Wnt/β-catenin pathway is recurrently activated in ∼20% of cancer and >50% of CIMP-high ACC (cBioPortal (Cerami et al., 2012; Gao et al., 2013), (Assie *et al*., 2014; Mohan *et al*., 2019; Zheng *et al*., 2016)). Efforts to target this pathway clinically have failed, due to life-limiting on-target toxicities in Wnt-dependent organs with rapid turnover (Kahn, 2014). Here, we discover that selection for active β-catenin in ACC is dependent on maintenance of a tissue specific program: SF1-dependent zF differentiation. Given the paucity of organs that require both programs for homeostatic renewal, this opens a large therapeutic window for targeting oncogenic β-catenin. Do alternative context-specific nuclear receptors bind β-catenin in other cancers? A paradigm geared towards tissue-specific disruption of oncogenic programs will be essential to combat cancers that rely on differentiation for growth and dissemination, like ACC.

Our studies illuminate how derailed epigenetic programs advantage cancer cells by maintaining a permissive chromatin environment for context-specific sustained proliferation. Here, we demonstrated through mechanistic and preclinical studies that dedifferentiation using epigenetic agents already FDA approved for other cancers (Morschhauser et al., 2020) represents a promising avenue for CIMP-high ACC. As we move forward to discover new classes of therapies to equip all patients fighting this disease, it will be crucial to understand how aberrant epigenetic patterning emerges in the adrenal cortex and forges subtype-specific routes to malignancy.

## ACKNOWLEDGEMENTS

This work is supported by the University of Michigan Rogel Cancer Center (grant to GDH, scholarship to DRM), the NIH through the University of Michigan’s Cancer Center Support Grant (5 P30 CA46592), FAPESP (2020/02988-1 to SKNM) and CNPq Universal (422140/2016-3 to SKNM). IF, CRL, AML, and GDH are/were supported by R01 DK062027 (grant to GDH). DRM, CRL, TE, RA, AML, and GDH are supported by the US Department of Defense (CA180750 to GDH and CA180751 to GDH, TE, and RJA). DRM is/was also supported by the University of Michigan Medical Scientist Training Program (T32 GM7863), the University of Michigan Doctoral Program in Cancer Biology, and The Drew O’Donoghue Fund. DTB is supported by R01 DK123694. JR was supported by American Heart Association grant 20CDA35320016. ALS and RO were supported by NIH grant R01 GM086610. ERL was supported by NIH grant R01 CA215981. AAA was supported by F31 CA247104. The authors would like to express their deepest gratitude to the patients, families, and advocates who have contributed essentially and immeasurably to advancing our understanding of ACC. The authors would also like to thank: Pat O’Day, for extraction of steroids and analysis of LC-MS/MS data. Amy Blinder, for assistance with human adrenal samples. Eric Wang, Trey Weaver, Emilia Pinto, for critical reading of this manuscript. Tom Wilson, Andy Muntean, Sundeep Kalantry, for reading components of this work and always providing thoughtful, frank feedback. Preeti Mohan, for her constructive input throughout project development. This paper was typeset with the bioRxiv word template by @Chrelli: www.github.com/chrelli/bioRxiv-word-template

## AUTHOR CONTRIBUTIONS

Conceptualization: AML, DRM, GDH; Data curation: DRM, AML; Formal Analysis: DRM, AML, SV, KSB; Funding acquisition: GDH, AML, TE, RJA, SKNM, DRM; Investigation: DRM, KSB, IF, CRL, JR, ALS, RO, DWL, TE, MQA, DD, JH-S, AAA, AW, SV, AML; Methodology: DRM, AML, KSB, IF, CRL, JR, AS, AA, AW, MCNZ, SV, SKNM, TJG; Project administration: DRM, AML, MCBVF, BMPM, MV, MCNZ; Resources: GDH, AML, DTB, MCBVF, SV, TJG, SKNM, RJA, WER, RO, ERL, BBM, ACL, LA; Software: AML, DRM; Supervision: AML, GDH, DTB; Validation: DRM, AML; Visualization: DRM, AML; Writing – original draft: DRM, AML; Writing – review & editing: All authors

## MATERIALS AND METHODS

### Resource availability

Original RNA-seq, methylation array, ChIP-seq, and ATAC-seq data have been deposited in GEO and will be publicly available as of the date of peer-reviewed publication.

### Experimental Model and Subject Details

#### Patient samples

All patient/donor samples used in this study were obtained with informed consent from the University of Michigan and Faculdade de Medicina da Universidade de Sao Paulo (FMUSP) as previously described (Mohan *et al*., 2019). Tissue microarrays (TMAs) containing benign, malignant, and metastatic adrenocortical tumors were developed and provided by FMUSP.

#### Cell culture

All cell lines were cultured under standard sterile conditions and maintained in a humidified tissue culture incubator with 5% CO2 at 37°C. NCI-H295R were obtained from ATCC and cultured in DMEM/F12 supplemented with 10% Nu serum, 1% ITS-X, and 1% penicillin/streptomycin. Y1 were obtained from ATCC and cultured in high glucose DMEM, supplemented with 2.5% fetal bovine serum, 7.5% horse serum, and 1% penicillin/streptomycin. ATC7L were a generous gift to our laboratory from A. Lefrançois-Martinez and A. Martinez (GReD, CNRS, Inserm, Université Clermont-Auvergne, Clermont-Ferrand, France) and cultured as previously described (Walczak *et al*., 2014), in DMEM/F12 supplemented with 2.5% fetal bovine serum, 2.5% horse serum, 1% ITS-X, and 1% penicillin/streptomycin. All cell lines were routinely screened (at every 3-5 passages and/or each experimental plating) for microbial contamination by DAPI staining and/or e-Myco Mycoplasma PCR Detection Kit (Bulldog Bio). Cell lines were discarded after 20-25 passages post-thaw, or when exponential growth was no longer evident (whichever came first).

#### Murine studies

All animal procedures were approved by Boston Children’s Hospital’s Institutional Animal Care and Use Committee. All mice were housed at Boston Children’s Hospital’s specific-pathogen free facility. The following adrenal-specific transgenic mice: (1) *AS*^*Cre/+*^, (2) p53*-*LOF (*AS*^*Cre/+*^:: *Trp53*^*flox/flox*^), PCre^AS/+^; (3) βcat-GOF (*AS-*^*Cre/+*^:: *Ctnnb*^*flox(ex3)/+*^), BCre^AS/+^; and (4) p53-LOF/βcat-GOF (*AS-*^*Cre/+*^:: *Trp53*^*flox/flox*^:: *Ctnnb*^*flox(ex3)/+*^), BPCre^AS/+^ have been previously described (Borges *et al*., 2020). Seven-to eight-week-old NOD.Cg-*Prkdc*^*scid*^ *Il2rg*^*tm1Wjl*^*/SzJ* (NSG) immunodeficient male mice were obtained from Jackson laboratories.

### Method Details

#### Nucleic acid extraction and quantification

Nucleic acid extraction from cells or patient samples was performed as described in (Mohan *et al*., 2019) with optional nuclease treatments, using the DNeasy Blood & Tissue Kit (Qiagen), using the RNeasy Plus Mini Kit (Qiagen), or using the TRI reagent (Sigma) with RNeasy kit (Qiagen) according to manufacturer’s instructions. Nucleic acid quantification was performed as described in (Mohan *et al*., 2019).

#### Targeted gene expression analysis

cDNA synthesis was performed on extracted mRNA as described in (Mohan *et al*., 2019) or according to manufacturer’s protocol using the iScript cDNA Synthesis Kit (Bio-Rad). SYBR qPCR was performed as described in (Mohan *et al*., 2019) using Power SYBR Green qPCR Mastermix (Invitrogen), and *Actb* (mouse) or *PPIB* (human) as the housekeeping gene. Targeted assessment of *G0S2* methylation was performed as previously described (Mohan *et al*., 2019). In patient samples, measurement of *BUB1B* expression, *GUSB* housekeeping gene expression, and *G0S2* methylation was performed and previously reported (Mohan *et al*., 2019). In this study, *EZH2* expression was measured by TaqMan Gene Expression Assays (Applied BioSciences/Thermo Fisher Scientific). Measurement of gene expression in BPCre^AS/+^ adrenals was performed using the QuantStudio 6 Flex Real-Time PCR System. In murine tissue samples, gene expression was measured by TaqMan Gene Expression Assays (Applied BioSciences/Thermo Fisher Scientific) with *Actb* as the housekeeping gene. Gene expression levels were calculated using the ΔC_t_ method as previously described (Mohan *et al*., 2019).

#### Brightfield IHC on TMAs

TMA of primary adrenocortical tumors from FMUSP was stained for EZH2 and H3K27me3 in triplicate. EZH2 staining was performed with Cell Signaling Technology, Cat. No. 5246S using standard IHC deparaffinization/hydration protocols and the NovoLink Max Polymer Detection System (Leica Biosystems) with modifications. EZH2 expression level was quantified on 0-4 scale based on % positive nuclei per section and averaged across two independent observers. H3K27me3 staining was performed as previously described (Bayliss *et al*., 2016). H3K27me3 levels were quantified by MATLAB as previously described (Bayliss *et al*., 2016).

#### Pharmacological experiments (not including NGS)

NCI-H295R cells were plated in 6 well plates, 24 well plates or 12 well plates. 18-24 hours after plating, media was changed for media containing EPZ-6438, EED226, GSK126, PRI-724, or vehicle (DMSO). The concentration of vehicle was kept constant for all wells in a given experiment. Media was replaced with fresh drug-containing media every 24 hours. After 96 hours of drug treatment, cells were harvested for RNA, protein or cell viability. Alternatively, cells were treated with PRI-724 and one of the following EZH2 inhibitors (EZH2i) at the IC-50 dose: EPZ-6438, EED226, GSK126 and harvested at 96 hours for cell viability. Alternatively, after 96 hours of EZH2i, cells were treated with forskolin for 48 hours, after which time media was harvested for steroid measurement and cells were harvested for protein or RNA. Alternatively, 18-24 hours after plating, media was changed for media containing forskolin, changed and replaced as above. After 48 hours of forskolin treatment, cells were harvested for RNA. Y1 or ATC7L were plated in 6 well plates. 18-24 hours after plating, media was changed to media containing forskolin or equivalent volume of vehicle (DMSO), media containing LiCl or equivalent volume of vehicle (water; % vehicle was kept constant for all doses), media containing 20% Wnt3a conditioned medium (CM) or 20% parental medium. Wnt3a CM or parental medium was derived from L cells as in (Shibamoto et al., 1998). Cells were harvested at the indicated timepoints. Gene expression from vehicle-treated cells in LiCl series were used for Y1/ATC7L comparison in **Supp Fig 3D**. Alternatively, pharmacological treatments on Y1 and ATC7L cell lines were performed similarly to those on NCI-H295R as above with the exception that Y1 and ATC7L cells were plated at a different density.

#### siRNA experiments

NCI-H295R were plated at 70-80% confluency in 6 well, 12 well or 24 well plates in antibiotic-free media. 18-24 hours after plating, media was changed for fresh antibiotic-free media containing 50 nM siRNA (Thermo Fisher) and transfection reagent (Mirus) prepared according to manufacturer instructions. Transfection was repeated 72 hours after first transfection and cells were harvested at 144 hours after first transfection for desired endpoint readouts.

#### Protein extraction and quantification

At endpoint, cells were washed with ice cold PBS and incubated on ice with ice cold whole-cell nuclear lysis buffer (50 mM Tris-HCl pH 8.1, 10 mM EDTA, 1% SDS in ultrapure H2O, adapted from (Drelon et al., 2016)), supplemented with protease inhibitors and phosphatase inhibitors according to manufacturer’s instructions. Cells were scraped off plate in lysis buffer and transferred to microcentrifuge tubes, and sonicated. After sonication, lysates were incubated on ice, centrifuged to remove insoluble debris, and supernatants collected in fresh tubes. Alternatively, cells were washed with PBS containing phosphatase inhibitors, scraped into microcentrifuge tubes, pelleted by centrifugation, frozen at -80°C, and later extracted for protein using complete whole-cell nuclear lysis buffer as stated above. Alternatively, cells were washed with PBS, incubated on ice with complete whole-cell nuclear lysis buffer, scraped into microcentrifuge tubes, frozen at -80°C and protein extraction was later resumed as stated above. Prior to downstream processing, lysates and protein standards were quantified by BCA (Thermo Scientific) in duplicate in 96 well plates according to manufacturer instructions. Absorbance was read at 562 nm using a VersaMax tunable microplate reader (Molecular Devices). Absorbance values of standard curve were used to determine sample concentration.

#### Cellular fractionation

Protein extracts were prepared using the ActiveMotif Nuclear Complex Co-IP Kit according to manufacturer’s protocol with supernatant from spin after hypotonic lysis saved as cytoplasmic fraction, and additional modifications.

#### SDS-PAGE and western blot

Samples were prepared by boiling at 95°C for 6 minutes in lysis buffer supplemented with 4x Laemmli Sample Buffer (Bio-Rad), with 355 mM 2-mercaptoethanol freshly added according to manufacturer’s instructions. Alternatively, samples were boiled at 95°C for 5 minutes in 2X reducing buffer (130 mM Tris pH 6.8, 4% SDS, 0.02% bromophenol blue, with 0.1 M DTT freshly added) with 16.7% glycerol. Samples and a size marker were loaded onto NuPage 4-12% Bis-Tris gels and run using the MOPS or MES buffer systems. Following SDS-PAGE, gels were coomassie stained or transferred onto a PVDF membrane using a traditional wet transfer system and NuPage Transfer Buffer supplemented with up to 20% methanol according to manufacturer instructions. Uniform transfer efficacy was verified by Coomassie staining of SDS-PAGE gel after transfer. Western blot was performed according to standard protocols. The following antibodies were used for western blot: EZH2 (Cell Signaling Technology, Cat. No. 5246S), SUZ12 (Cell Signaling Technology, Cat. No. 3737), SUZ12 (R&D, Cat. No. MAB4184), active β-catenin (Cell Signaling Technology, Cat. No. 8814), β-catenin (Invitrogen, Cat. No. MA1-2001), H3K27me3 (EMD Millipore, Cat. No. 07-449), H3K27ac (ActiveMotif, Cat. No. 39133), SF1 (custom in-house antibody, RRID AB_2716716), SF1 (Invitrogen, Cat. No. 434200/N1665), Lef1 (Cell Signaling Technology, Cat. No. 2230), β-actin (Sigma-Aldrich, Cat. No. A-5441).

#### Coomassie staining

Coomassie staining was performed using Imperial Protein Stain (Thermo) according to manufacturer instructions. After overnight destaining, gels were imaged using the LI-COR Odyssey imaging system protein gel setting at 700 nm.

#### Size-exclusion chromatography

Nuclear lysates were prepared from NCI-H295R or Y1 cell lines using ActiveMotif Nuclear Complex Co-IP Kit. Sample was clarified via centrifugation. Supernatant was subjected to size exclusion chromatography on a Superose 6 column (GE Healthcare) equilibrated in a buffer of 20 mM HEPES (pH 7.9), 75 mM NaCl, and 1 mM DTT. Eluents spanning elution volumes 10 mL to 45 mL were collected and fractionated in 0.5 mL fractions. Approximate protein size of fractions was estimated using an empirical calibration curve for the Superose 6 column. Fractions were analyzed by SDS-PAGE, Coomassie staining, and western blot.

#### Nuclear complex immunoprecipitation (co-IP)

Nuclear co-IP was performed using the ActiveMotif Nuclear Complex Co-IP Kit and Protein G Agarose Columns according to manufacturer’s protocol with modifications. Nuclear lysate concentration was quantified by BCA prior to setting up co-IP. Any of the following antibodies were used for co-IP: EZH2 (Cell Signaling Technology, Cat. No. 5246S), SUZ12 (Cell Signaling Technology, Cat. No. 3737), SUZ12 (Bethyl, Cat. No. A302-407A), active β-catenin (Cell Signaling Technology, Cat. No. 8814), SF1 (custom in-house antibody, RRID AB_2716716), negative control IgG (EMD Millipore, Cat. No. 12-370). Co-IPs were evaluated by mass spectrometry (IP-MS), or were eluted and evaluated by Coomassie staining and/or western blot.

#### Immunoprecipitation/mass spectrometry (IP-MS)

NCI-H295R cells were cultured as described and plated in 10 cm dishes. To minimize biological variability, 3 ∼80% confluent plates of 3 independent passages of exponentially growing cells (total 9 plates) were harvested and pooled for nuclear co-IP using the ActiveMotif Nuclear Complex Co-IP Kit and Protein G Agarose Columns. Nuclear co-IP was performed as described using aforementioned antibodies against EZH2, active β-catenin, SF1, DNMT1, or negative control IgG with the following modifications: Co-IPs were not eluted. Instead, after final wash, samples were rinsed in PBS, and Protein G columns were stored in new 1.5 mL microcentrifuge tubes and frozen at -80°C. Samples were delivered on dry ice to MS Bioworks, Ann Arbor, MI for mass spectrometry analysis using the IP-works platform.

IP-MS data were searched using a local copy of Mascot (Matrix Science) with the following parameters: Enzyme – Trypsin/P, Database – SwissProt Human (concatenated forward and reverse plus common contaminants), Fixed modification – Carbamidomethyl (C), Variable modifications – Oxidation (M), Acetyl (N-term), Pyro-Glu (N-term Q), Deamidation (N/Q), Mass values – Monoisotopic, Peptide Mass Tolerance – 10 ppm, Fragment Mass Tolerance – 0.02 Da, Max Missed Cleavages – 2. Mascot DAT files were parsed into Scaffold (Proteome Software) for validation, filtering and to create a non-redundant list per sample. Data were filtered using at 1% protein and peptide FDR and requiring at least two unique peptides per protein. Known contaminants and reverse hits were excluded from downstream analysis.

In this study, a protein was considered an interactor if it met the following criteria: ≥ 5 spectral counts in target co-IP and either not detected in IgG co-IP or at least 2-fold enrichment over IgG in target co-IP based on dividing spectral count values. Prior to submission of co-IPs for IP-MS, co-IP efficacy was verified by Coomassie staining and evaluation of *bona fide* protein interactions reported in the literature by western blot (e.g. EZH2/SUZ12, DNMT1/PCNA). The 2-fold cutoff for enrichment was determined based on the minimum threshold required to capture *bona fide* protein interactors (supported by the literature).

#### Viability assays and calculations

Viability was measured using alamar-Blue (Invitrogen) according to manufacturer’s instructions. Absorbance was measured using VersaMax tunable microplate reader (Molecular Devices) at 570 nm using 600 nm as a reference wavelength. % viability (%V) was calculated as follows. For vehicle or each treatment concentration, AR=(ε_OX,600_)(A_570_–B_570_)–(ε_OX,570_)(A_600_–B_600_), where ε_OX,600_=117,216, ε_OX,570_=80,586, A_570_=absorbance of treatment well at 570 nm, B_570_=absorbance of blank well (well containing no cells, only media and alamar-Blue) at 570 nm, A_600_=absorbance of treatment well at 600 nm, and B_600_=absorbance of blank well at 600 nm. %V=100*AR_treatment_/AR_vehicle_. Data were fitted to a 4PL sigmoidal curve using GraphPad and IC-50 was determined from interpolation of the curve to identify the concentration at 50% viability. To evaluate drug interactions, a simplified coefficient of drug interaction (CDI) was calculated as described (Pham et al., 2019). CDI>1 signifies antagonism, CDI=1 signifies additivity, and CDI<1 signifies synergy. Negative CDI values are secondary to fluctuations of absorbance around the level of the blank when no viable cells remain in a well, and are consistent with CDI<1 for a drug that induces loss of viability. An alternative strategy would be to set the values of these conditions to 0.

#### 2D Clonogenicity

At experiment endpoint, cells were washed with warm PBS, trypsinized, and viable cells (identified by Trypan blue exclusion) were counted using a hemocytometer. 1,000 viable cells/well were plated in 6 well plates in standard media. Cells were maintained under standard conditions with no media changes. 4 weeks after plating, plates were washed with PBS and fixed with 4% PFA. Plates were washed with water, stained with crystal violet staining solution (0.1% crystal violet in 5% ethanol in water), washed with water, and inverted over a paper towel to dry overnight. The next day, plates were imaged using the LI-COR Odyssey imaging system using the microplate setting at 700 nm. Colonies were counted using the “Analyze Particles” tool in Fiji (Schindelin et al., 2012).

#### RNA-seq

NCI-H295R were plated in 6 well plates and treated for 96 hours as above with EZH2i at the IC-50 dose, CBPi at the IC-50 dose, or vehicle (equivalent volume of DMSO, these are “baseline”) in 3 biological replicates. Alternatively, NCI-H295R were plated in 6 well plates and treated with equivalent volume of vehicle (DMSO) for 48 hours and then treated with forskolin for 48 hours as above in 3 biological replicates. mRNA and genomic DNA were extracted at endpoint for RNA-seq and 850k array profiling, respectively. RNA-seq was performed by the University of Michigan Advanced Genomics Core. Libraries were prepared from total mRNA extracted from NCI-H295R according to standard Illumina protocols. 50 bp paired-end reads were generated in the Illumina NovaSeq-6000 platform (S1 100 cycle) at an output to ensure ∼50 million reads/sample. Reads were aligned to the hg38 assembly of the human genome using STAR (Dobin et al., 2013). Gene expression was quantified by featureCounts (Liao et al., 2014). Quality control metrics were generated by RNA-SeQC (DeLuca et al., 2012). Count data was normalized using TMM normalization from edgeR (McCarthy et al., 2012; Robinson et al., 2010); logCPM values were generated using the voom-WithQualityWeights function from limma (Ritchie et al., 2015). FPKM values were calculated using edgeR (McCarthy *et al*., 2012; Robinson *et al*., 2010). Genes were ranked by FPKM to determine percentile of expression. GSVA (Hänzelmann *et al*., 2013) was used to calculate zF differentiation, cell cycle, and Wnt scores per class with genes as described in methods section detailing analysis of ACC-TCGA data. BAM files for Y1 RNA-seq were generated according to the same pipeline from fastq files downloaded from DDBJ/EMBL/GenBank DRA000853 aligned to mm10.

#### ATAC-seq

NCI-H295R were plated in 6 well plates and treated for 96 hours as above with EZH2i at the IC-50 dose, CBPi at the IC-50 dose, or vehicle (equivalent volume of DMSO, these are “baseline”) in 2 biological replicates. Alternatively, NCI-H295R were plated in 6 well plates and treated with equivalent volume of vehicle (DMSO) for 48 hours and then treated with forskolin for 48 hours as above in 2 biological replicates. At endpoint, cells were harvested for ATAC-seq. ATAC-seq was performed as previously described (Corces et al., 2017) with modifications. Tn5 transposase-tagged DNA was amplified by PCR and purified using AMPure XP magnetic beads. ATAC-seq reads were sequenced 2×150 bp using a NovaSeq 6000. Sequencing of ATAC-seq libraries was performed by the University of Michigan Advanced Genomics Core. We used bowtie2 (Langmead and Salzberg, 2012) to align the reads to the hg38 version of the human genome. Reads overlapping with blacklisted regions (defined by ENCODE (Consortium, 2012; Davis *et al*., 2018)), and reads with a mapping score < 20 were filtered. For ATAC-seq peak calling, we used genrich (Gaspar, 2018). For differential peak calling we used diffbind (Ross-Innes et al., 2012; Stark and Brown, 2011). Integration of RNA-seq and ATAC-seq data was performed using diffTF (Berest *et al*., 2019). BigWig files were generated from merged replicates using deepTools 2.0 (Ramírez et al., 2016) with RPGC normalization, and visualized using the JBR browser (Shpynov et al., 2021) or BigwigTrack from Signac (Stuart et al., 2021).

#### ChIP-seq

NCI-H295R were plated in standard culture medium in 10 cm dishes. To best account for biological variability, this experiment was performed using 24 plates of cells total, where 16 plates were reserved for EZH2i administration at the IC-50 dose and 8 plates were reserved for vehicle administration (equivalent volume of DMSO, these are considered “baseline” NCI-H295R). 24 hours after plating and daily thereafter, media was changed for media containing IC-50 EZH2i or vehicle. After approximately 96 hours of drug administration (120 hours post-plating), cells were fixed and harvested for ChIP-seq with *Drosophila melanogaster* histone spike-in for all epitopes according to Active Motif’s Epigenetic Services ChIP Cell Fixation protocol.

Briefly, chromatin was isolated by the addition of lysis buffer, followed by disruption with a Dounce homogenizer. Lysates were sonicated and DNA sheared to an average length of 300-500 bp. Genomic DNA (Input) was prepared by treating aliquots of chromatin (pooled from all submitted samples) with RNase, proteinase K and heat for de-crosslinking, followed by ethanol precipitation. Pellets were resuspended and the resulting DNA was quantified on a NanoDrop spectrophotometer. Extrapolation to the original chromatin volume allowed quantitation of the total chromatin yield.

An aliquot of chromatin was precleared with protein A agarose beads (Invitrogen). Genomic DNA regions of interest were immunoprecipitated using the following antibodies: EZH2 (Active Motif, Cat. No. 39901), H3K27me3 (ActiveMotif, Cat. No. 39155), H3K27ac (ActiveMotif, Cat. No. 39133), SF1 (EMD Millipore, Cat. No. 07-618) and β-catenin (Invitrogen, Cat. No. 71-2700). Complexes were washed, eluted, and subjected to RNase and proteinase K treatment and crosslink reversal. ChIP DNA was purified by phenol-chloroform extraction and ethanol precipitation. Quantitative PCR (qPCR) reactions to verify ChIP efficacy were carried out in triplicate on specific positive control genomic regions using SYBR Green Supermix (Bio-Rad).

Illumina sequencing libraries were prepared from the ChIP and Input DNAs by the standard consecutive enzymatic steps of end-polishing, dA-addition, and adaptor ligation. Steps were performed on an automated system (Apollo 342, Wafergen Biosystems/Takara). After a final PCR amplification step, the resulting DNA libraries were quantified and sequenced on Illumina’s NextSeq 500 (75 nt reads, single end).

We used bowtie2 (Langmead and Salzberg, 2012) to align the reads to the hg38 version of the human genome and the dm10 version of the *Drosophila melanogaster* genome. Reads overlapping with blacklisted regions (defined by ENCODE), and reads with a mapping score < 20 were filtered. BAM files were down-sampled to account for *Drosophila* spike-in, enabling quantitative comparison of epitopes accounting for net chromatin recruitment across conditions. We performed ChIP-seq peak calling using SPAN and the JBR browser (Shpynov *et al*., 2021). We used the annotatePeak function from the R/Bioconductor package ChipSeeker (Yu et al., 2015) to annotate the peaks according to distance to TSS and overlapping features. For motif enrichment analysis we used HOMER (Heinz *et al*., 2010). We used bedtools (Quinlan and Hall, 2010) for identifying intersections of interest across different epitopes. BigWig files were generating using deepTools 2.0 (Ramírez et al., 2016) with RPGC normalization, and visualized using the JBR browser (Shpynov et al., 2021) or BigwigTrack from Signac (Stuart et al., 2021). For visualization and quantification of the intensity of the signal across peaks of interest we used deepTools 2.0 (Ramírez et al., 2016). We identified super-enhancers using ROSE (Lovén *et al*., 2013; Whyte *et al*., 2013).

#### Methylation arrays

Extracted gDNA from patient samples was subject to 850k array profiling performed by the University of Michigan Advanced Genomics Core. Extracted gDNA from EZH2i-/vehicle-treated NCI-H295R in biological triplicate was submitted to Diagenode Epigenomic Services, Denville, NJ, USA for 850k array profiling. MethylAid (van Iterson et al., 2014) was used on 850k array data to assess quality control, and minfi (Aryee et al., 2014) was used to perform functional normalization, and generate table of beta values. Table was filtered for probe sets of interest. PRC2 target probes were probes that fall in the promoters of genes included in the BENPORATH_PRC2_TARGETS set deposited in GSEA (Ben-Porath *et al*., 2008; Liberzon et al., 2011; Subramanian *et al*., 2005). Copy number inference was performed using conumee (Hovestadt and Zapatka, 2020). Limma (Ritchie *et al*., 2015) was used for statistical analysis to assess probe-specific differential methylation.

#### Brightfield IHC (excluding H3K27me3 on allografts)

IHC was performed using the VECTASTAIN Elite ABC-HRP Kit, Peroxidase (Rabbit IgG) +/-the M.O.M. (Mouse on Mouse) Immunodetection Kit – Basic (Vector Laboratories, Cat. No. BMK-2022) according to manufacturer’s instructions with modifications. Samples were stained using antibodies against SF1 (RRID AB_2716716), β-catenin (BD Biosciences, Cat. No. 610154), EZH2 (Cell Signaling Technology, Cat. No. 5246S) or Ki67 (Thermo Fisher MA5-14520). 3,3’-Diaminobenzidine (DAB) served as the peroxidase substrate and were used according to manufacturer’s instructions with same development time for all samples. Slides were counterstained with Gills #1 Hematoxylin. IHC DAB signal was quantified on 10 40x images per mouse using a homemade macro in Fiji (Schindelin *et al*., 2012). Briefly, for Ki67 signal: Images were deconvoluted using H and DAB vectors, thresholded at (0, 225) and (0, 210) respectively, converted to mask, watershedded, and Analyze Particles function was used to count nuclei and DAB staining (minimum particle size 400). DAB/nuclear signal was computed per image and averaged to yield Ki67 (%) value for each sample. Briefly, for SF1 signal: same as for Ki67 except images were thresholded at (0, 235) and (0, 180), respectively.

#### Brightfield H3K27me3 allograft IHC

Studies were performed as previously described (Chung et al., 2020; Panwalkar et al., 2017) using anti-H3K27me3 (07-449, Millipore-Sigma). Each section was scanned using an AperioScanscope Scanner (Aperio Vista) and visualized using the AperioImageScope software program. Three random JPEG images from each section at 40X magnification were captured by an individual blinded to the experimental setup and quantified with an automated analysis program with MATLAB’s image processing toolbox (Chung *et al*., 2020; Panwalkar *et al*., 2017). We used algorithms based on K-Means Clustering, color segmentation with RGB color differentiation, and background-fore-ground separation with Otsu’s thresholding (Panwalkar *et al*., 2017). The numbers of extracted pixels were then multiplied by their average intensity for each image and was represented as pixel units.

#### Immunofluorescence

Adrenals were fixed in 4% paraformaldehyde (PFA) for 2 hours (normal tissue, 24.3 week female in **Figure 5B**) or overnight (tumor, from 41.1 week male in **Figure 5B**), embedded in paraffin and cut into 5-μm sections, and immunofluorescence was performed using standard protocols as previously described with DAPI nuclear staining. Primary antibodies: β-catenin (BD Biosciences, Cat. No. 610153), SF1 (Thermo Fisher, Cat. No. 434200), H3K27me3 (Cell Signaling Technology, Cat. No. 9733) and EZH2 (Cell Signaling Technology, Cat. No. 5246). Secondary antibodies: Alexa Fluor 647-conjugated goat anti-rabbit IgG (Invitrogen), Alexa Fluor 594-conjugated goat anti-mouse IgG (Invitrogen). Images were acquired using a Nikon upright Eclipse 90i microscope and entire images were adjusted for brightness and contrast in ImageJ (Schneider et al., 2012).

#### Proximity Ligation Assay (PLA)

PLA was performed using the Duolink Detection Reagents Brightfield Kit (Sigma-Aldrich, Cat. No. DUO92012) and associated wash buffers (Sigma-Aldrich, Cat. No. DUO82047-4L) and PLA probes according to manufacturer protocol with modifications to blocking and primary antibody incubation steps. Counterstain step was skipped because counterstain masked signal from nuclear epitopes. The following antibodies were used for PLA in murine dysplasia to carcinoma sequence: EZH2 (Cell Signaling Technology, Cat. No. 5246S), β-catenin (BD Biosciences, Cat. No. 610154), SF1 (custom in-house antibody, RRID AB_2716716), SF1 (Invitrogen, Cat. No. 434200/N1665). They were used at the following combinations: EZH2/β-catenin to detect EZH2/β-catenin interactions, SF1 (Rb)/β-catenin to detect SF1/β-catenin interactions, SF1 (Rb)/SF1 (Mmu) to detect SF1+ cells in metastatic lesions, or no antibodies (No Ab) as a negative control. The following antibodies were used for PLA in human tissue including FMUSP TMAs of primary and metastatic adrenocortical tumors in triplicate: EZH2 (Cell Signaling Technology, Cat. No. 5246S), β-catenin (BD Biosciences, Cat. No. 610154), SF1 (custom in-house antibody, RRID AB_2716716). They were used at the following combinations: SF1/β-catenin to detect SF1/β-catenin interactions or EZH2/β-catenin to detect EZH2/β-catenin interactions. Extensive optimization was performed on mouse and human tissue prior to arriving at optimal blocking conditions and antibody concentrations, including combining concentrations of antibodies at different combinations, and testing out positive control and no antibody/single primary antibody incubations. For example, as a positive control (detecting β-catenin) in a small cohort of samples and during PLA optimization, Active β-catenin, Cell Signaling Technology, Cat. No. 8814 was paired with varying concentrations of β-catenin (BD Biosciences, Cat. No. 610154) or β-catenin (Invitrogen, Cat. No. MA1-2001).

Quantification of PLA signal: TMA - PLA signal was quantified on 3 20x images per replicate using a homemade macro in Fiji (Schindelin *et al*., 2012). Nuclei were quantified using the greater of the two values computed using Method 1 or 2. Method 1 - Images were deconvoluted using H&E vectors. Pink signal was thresholded at (0, 230), converted to mask, watershedded, and Analyze Particles function was used to count nuclei (minimum size 350). Method 2 - Images were deconvoluted using H&E 2 vectors. Pink signal was thresholded at (0, 230), converted to mask, watershedded, and Analyze Particles function was used to count nuclei (minimum size 350). PLA dots were quantified as described in Method 2 with the following modifications: Blue signal was thresholded at (0, 151), and particles with size 5-100 were counted with Analyze Particles. PLA/nuclear signal was computed per image and averaged to yield normalized PLA signal for each sample. Allografts - PLA signal from allografts was quantified as described except that it was quantified on 10 40x images per mouse.

#### LC-MS/MS

Steroids were extracted from culture medium and quantitated using liquid chromatography and tandem mass spectrometry as previously described (Wright et al., 2020). Protein was harvested from corresponding cells and quantified as above. Steroid measurements were normalized to total sample protein.

#### BCH-ACC3A allograft experiments

The mouse ACC cell line BCH-ACC3A was derived from a BPCre^AS/+^ male mouse harboring a large tumor (538 mg) with metastasis in the lung. Briefly, primary ACC from the mouse was dissociated as single cells using the mouse Tumor Dissociation Kit (Miltenyi Biotec). Cells were then serially transplanted into NSG mice for approximately 6 months. Cells were then cultivated *in vitro* in DMEM/F12 supplemented with 2% Nu serum, 1% ITS, 2 mM L-glutamine and 1% penicillin/streptomycin. Cells (5 × 10^4^) were resuspended in PBS mixed with Matrigel Basement Membrane Matrix (BD Biosciences) at 1:1 ratio and injected into subcutaneous tissue in the NSG mouse flank. Tumor length (L) and width (W) were measured with a digital caliper, and tumor volumes were calculated using the formula L*W^2^*(π/6) where L<W. When subcutaneous tumors achieved volume ∼100 mm^3^, mice were randomized to treatment with EPZ-6438 or equivalent volume of vehicle daily by oral gavage. Each dose was delivered in a volume of 10μL/g mouse.

### Quantification and Statistical Analysis

For all studies with a bench component, data processing and analysis are described above. Statistical information including number of replicates and statistical tests performed to compare experimental groups are described in methods above, figures and figure legends. P-value or adjusted p-value < 0.05 was considered significant for all analyses. Statistical analyses were performed using GraphPad Prism or R (Team, 2016). Pheatmap (RRID:SCR_016418) was used to perform unsupervised hierarchical clustering.

#### Analysis of ACC-TCGA

Differential gene expression analysis and retrieval of expression data from ACC-TCGA data was performed as described in (Mohan *et al*., 2019). Genes comprising zF differentiation score were identified using BioGPS and expert curation. Briefly, we selected transcripts that were highly differentially expressed in adrenal cortex compared to other organs as in (Zheng *et al*., 2016). This list was further curated to include genes that participate in steroid production, and augmented to include genes known to participate in steroidogenesis and adrenal differentiation. Given the importance of tissue of origin in defining program activation, we used independent component analysis (ICA) (Biton *et al*., 2014) across ACC-TCGA samples to identify modules of coregulated cell cycle and Wnt/β-catenin-regulated genes. The final lists of genes comprising the zF differentiation, Wnt, and cell cycle scores each included 25 genes.

We used GSVA (Hänzelmann *et al*., 2013) to calculate zF differentiation, cell cycle, and Wnt scores in ACC-TCGA samples. Final zF differentiation, cell cycle, and Wnt scores were validated on the basis of significantly higher expression in tumors with cortisol production, cell cycle pathway alterations (driver alterations in *TP53, RB1, CDKN2A, CCNE1, CDK4*), or Wnt pathway alterations (driver alterations in *CTNNB1, APC* or *ZNRF3*), respectively (**Supp Fig 1A-C**). Cell cycle and Wnt scores were further validated on the basis of significant correlation with *bona fide* cell cycle or Wnt pathway target genes (**Supp Fig 1D-E**). Correlograms were plotted using GGally. Regions targeted for differential methylation in CIMP-high vs. non-CIMP-high ACC were identified using DMRcate and published in (Mohan *et al*., 2019). Annotated significant (FDR-corrected p-value < 0.05) DMRcate regions were filtered to include only regions overlapping with promoters. Regions were ranked by descending mean difference in DNA methylation (beta fold change), and top 2084 gene identifiers (1994 NCBI Entrez Gene IDs, the maximum) were evaluated for enrichment with curated collections in GSEA (Mootha *et al*., 2003; Subramanian *et al*., 2005). ACC-TCGA ATAC-seq BigWig files were downloaded from TCGA Genomic Data Commons (NCI, 2005-2018), and signal at SF1/β-catenin binding sites was measured using multiBigwig-Summary in deepTools (Ramírez *et al*., 2016).

#### Analysis of microarray data from microdissected adrenals

Raw microarray data was downloaded from GEO (GSE68889) and analyzed as in (Finco *et al*., 2018) with RMA normalization (Irizarry et al., 2003).

#### ENCODE

The following datasets were downloaded from the ENCODE portal (Consortium, 2012; Davis *et al*., 2018) for analysis in this study. Hg38 H3K27ac ChIP-seq libraries from adrenal tissue, which was analyzed to generate BigWig files and peak calls per pipeline above: ENCLB278KZ2X, ENCLB060VHC, ENCLB923ALY, ENCLB245VYO, ENCLB872HUE, ENCLB191XGX, ENCLB281DGJ, ENCLB670CFC, ENCLB382MNY, ENCLB198CGW, ENCLB172SDF, ENCLB735YID. Hg38 H3K27me3 ChIP-seq BigWigs +/-peak calls from embryonic stem cells and adrenal tissue: H1 - ENCFF050CUG, ENCFF671NVD; H1 MSC - ENCFF639HFP, ENCFF998IEU; H1 NSC - ENCFF202US, ENCFF495GNE; H7 - ENCFF154IMT, ENCFF662NTZ; H9 - ENCFF342DVJ, ENCFF680AKW; HUES6 - ENCFF456EQK, ENCFF725EMZ; HUES48 - ENCFF009PDQ, ENCFF612HRW; HUES64 - ENCFF015TOT, ENCFF441TOI; UCSF-4 ENCFF218TIK, ENCFF342EGR; Fetal adrenal - ENCFF250ASJ, ENCFF495YGS; Adult adrenal - ENCFF019XYA.

#### Analysis of single-cell RNA-seq (scRNA-seq)

Gene expression matrices of human fetal, neonatal and adult adrenal scRNA-seq data (Han *et al*., 2020) were downloaded from GEO (GSE134355) and filtered, normalized, batch-corrected, integrated, scaled, and UMAP clustered using Seurat with CCA integration algorithm (Stuart et al., 2019). Gene expression was scaled and regressed against mitochondrial content and total number of transcripts/cell. Non-adrenocortical/capsular cells (e.g. immune cells, medulla cells) were excluded by serial rounds of UMAP clustering and cluster marker identification. Cluster marker identification also enabled assignment of single cell clusters to known adrenocortical/capsular cell populations. Adrenocortical/capsular cells were subject to pseudotime trajectory analysis using Monocle 3 (Cao et al., 2019; Qiu et al., 2017a; Qiu et al., 2017b; Trapnell et al., 2014), with UMAP embeddings from Seurat with origin set per figure legend.

#### Analysis of single-cell ATAC-seq (scATAC-seq)

ScATAC-seq was processed and plotted using Signac (Stuart *et al*., 2021) unless stated otherwise. Filtered hg38-aligned adult adrenal scATAC-seq peak matrices and fragment files were downloaded from GEO (GSM5047828) (Zhang *et al*., 2021). Adult adrenal scATAC-seq was processed to generate peak calls, converted to hg19 with rtracklayer (Lawrence et al., 2009), and then normalized, LSI dimension reduced, and UMAP clustered. Filtered hg19-aligned fetal adrenal scATAC-seq peak matrices and fragment files were downloaded from Descartes Human Chromatin Accessibility During Development Atlas (Domcke *et al*., 2020) (descartes.brotmanbaty.org) and processed as above (omitting rtracklayer step). Fetal adrenal scATAC-seq cells were downsampled to the same number of cells as in adult adrenal scATAC-seq. Fetal and adult adrenal peak sets were merged to generate a consensus peak set. Datasets were batch-corrected, integrated, normalized, LSI dimension reduced and UMAP clustered. Gene activity matrices were generated, and non-adrenocortical/capsular cells were excluded by serial rounds of UMAP clustering and cluster marker identification. Final clusters were compared to reference markers identified by scRNA-seq to make cluster assignments. Cells which could not be definitively classified were grouped together and unlabeled as shown in figure.

#### Identification of enhancer/gene links

Promoter-other contact tables derived from adrenal promoter capture Hi-C (pcHi-C) were downloaded from GEO (GSE86189) (Jung *et al*., 2019). An active enhancer is defined as a distal CRE overlapping with an H3K27ac peak. Putative active enhancers of a given gene were identified by overlapping adrenal pcHi-C contact tables with H3K27ac peaks identified from H3K27ac ChIP-seq on baseline NCI-H295R. To identify genes putatively regulated by active SF1/β-catenin enhancers, the set of active enhancers was then overlapped with the consensus SF1/β-catenin peak set. Otherwise, active enhancers were manually inspected for epitope of interest. Overlaps between regions were computed using bedtools (Quinlan and Hall, 2010).

---

**Supplementary Figure 1:**
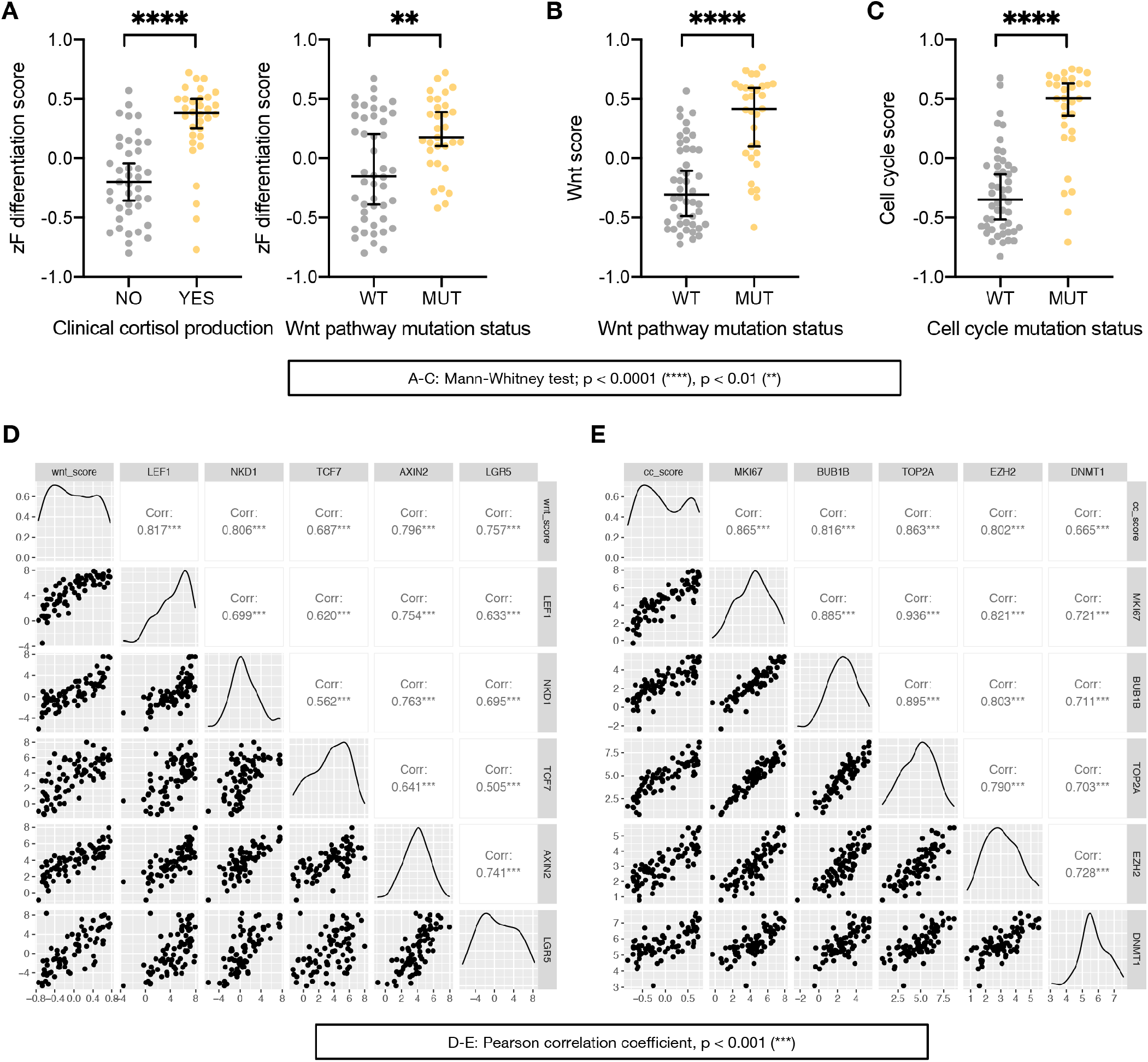
Validation of ACC-TCGA scores, related to Figure 1. **(A-C)** Genes comprising zF differentiation score, Wnt score, and cell cycle score were identified and calculated as described in **Materials and Methods**. Shown in **(A)**, zF differentiation score is significantly higher in ACC-TCGA tumors with clinical cortisol production, left, and Wnt pathway alterations (WT=Wnt wild type, MUT=driver alterations in *CTNNB1, APC*, or *ZNRF3*), right. Shown in **(B)**, Wnt score is significantly higher in ACC-TCGA tumors with Wnt pathway alterations. Shown in **(C)**, cell cycle score is significantly higher in ACC-TCGA tumors with cell cycle alterations (WT=Cell cycle wild type, MUT=driver alterations in *TP53, RB1, CDKN2A, CCNE1, CDK4*). **(D)** Across ACC-TCGA, Wnt score exhibits strong correlation with expression of *bona fide* canonical Wnt target genes. **(E)** Across ACC-TCGA, Cell cycle score exhibits strong correlation with expression of *bona fide* cell cycle markers (*MKI67, BUB1B, TOP2A*) and epigenetic modifiers (*EZH2, DNMT1*).

**Supplementary Figure 2:**
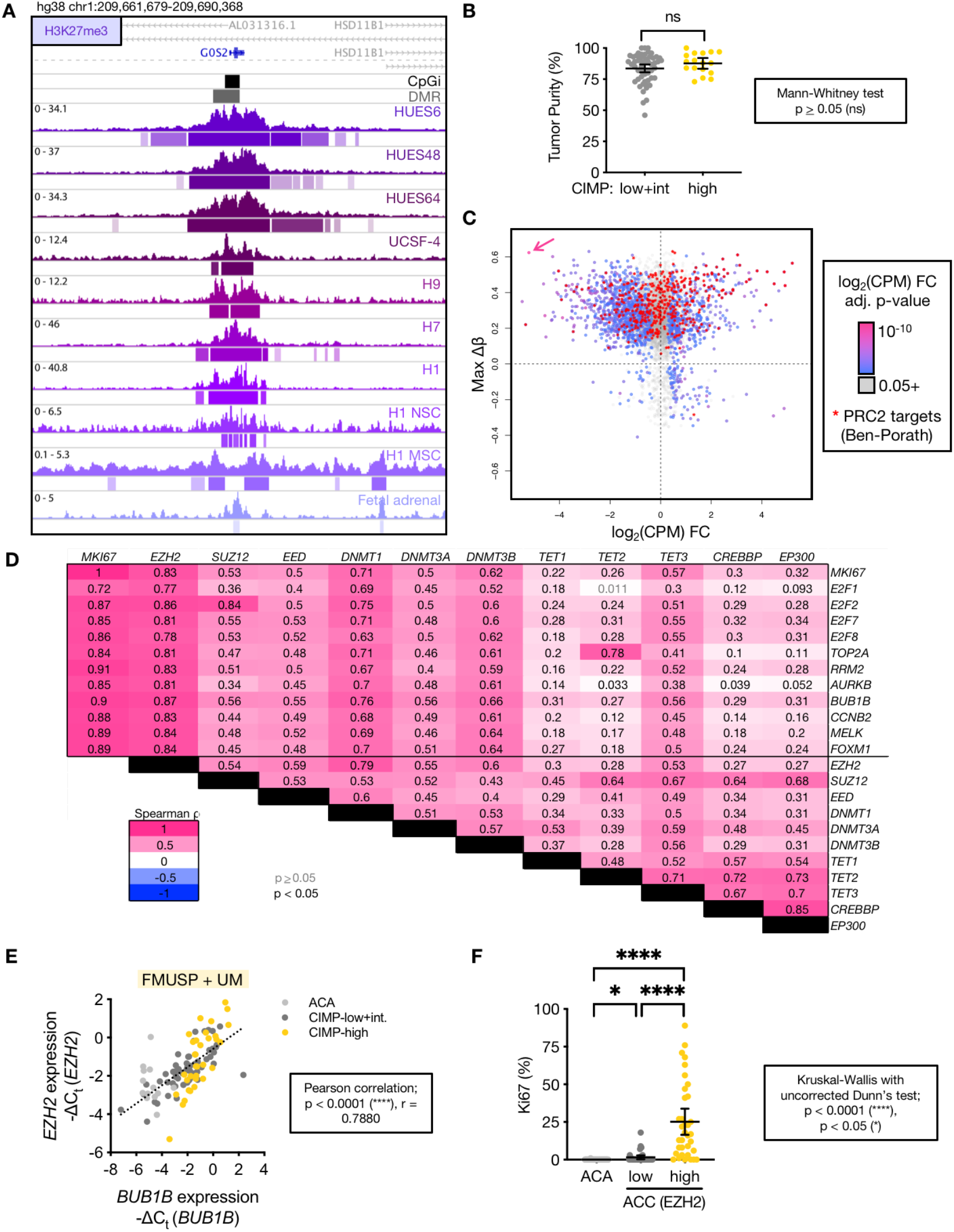
Related to Figure 1. **(A)** Example H3K27me3 ChIP-seq signal at the *G0S2* locus in human embryonic stem cells (ESC, includes HUES6, HUES48, UCSF-4, H9, H7, H1), ESC-derived neuronal stem cells (H1 NSC), ESC-derived mesenchymal stem cells (H1 MSC), and the fetal adrenal. Location of *G0S2* CpG island in black (top track), with region targeted for methylation in CIMP-high ACC depicted in grey (DMR, second track). Peak calls are depicted by bars below each track. Hg38-aligned ChIP-seq data and peak calls downloaded from ENCODE (Consortium, 2012; Davis *et al*., 2018). **(B)** Tumor purity across non-CIMP-high and CIMP-high ACC from ACC-TCGA (Zheng *et al*., 2016). Line at mean with 95% confidence interval (CI). **(C)** Scatterplot depicts max change in DNA methylation (max Δβ) between CIMP-high and non-CIMP-high ACC at promoter DMRcate regions (queried by GSEA in **Figure 1C** and originally reported in (Mohan *et al*., 2019)) versus log2 of the fold change in gene expression (measured by counts per million, CPM) between CIMP-high and non-CIMP-high ACC, color-coded by the p-value for the fold change in gene expression. PRC2 targets (from BENPORATH_PRC2_TARGETS set (Ben-Porath et al., 2008) deposited in GSEA) are indicated by red stars. *G0S2*, silenced by methylation in CIMP-high ACC, is indicated by the arrow. **(D)** Pan-cancer correlation matrix depicting relationship between cell cycle genes (*MKI67, E2F1, E2F2, E2F7, E2F8, TOP2A, RRM2, AURKB, BUB1B, CCNB2, MELK, FOXM1*) and genes encoding epigenetic modifiers (*EZH2, SUZ12, EED, DNMT1, DNMT3A, DNMT3B, TET1, TET2, TET3, CREBBP, EP300*) across TCGA. Expression data with Spearman ρ and p-value for each pair-wise comparison were mined and computed with GEPIA (Tang et al., 2017). **(E)** *EZH2* expression vs. expression of cell cycle-dependent gene *BUB1B* across adrenocortical tumors measured by qPCR in our independent cohort. Cohort with *BUB1B* expression data originally published in (Mohan *et al*., 2019). **(F)** Ki67 proliferation index (% nuclei Ki67+) across ACA and primary ACC stratified by EZH2 expression in TMA. EZH2 low=ACC with below median EZH2 expression (≤ 25% nuclei EZH2+), EZH2 high=ACC with above median EZH2 expression (> 25% nuclei EZH2+). ACA, n=74; EZH2 low ACC, n=37; EZH2 high ACC, n=35.

**Supplementary Figure 3:**
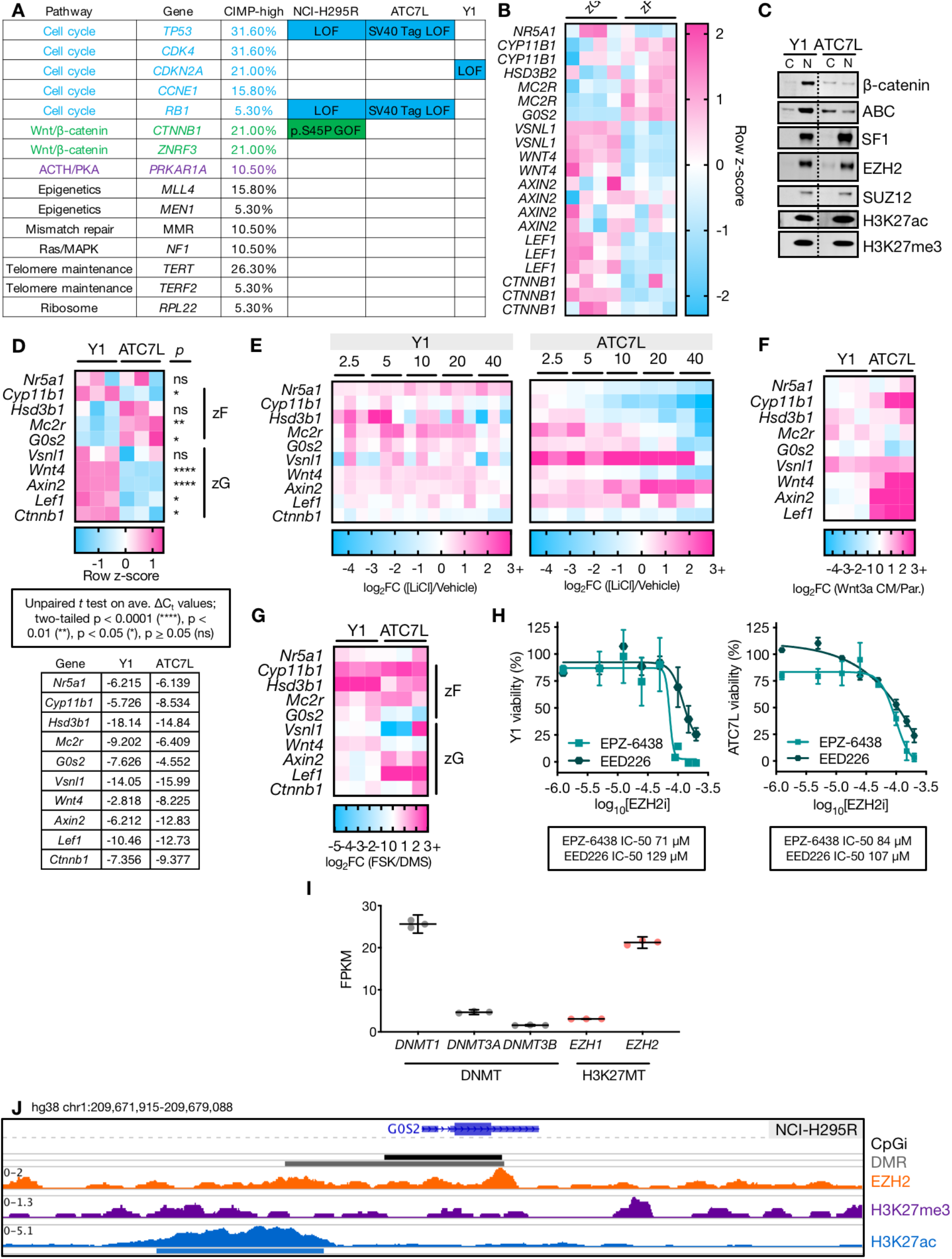
Related to Figure 2. **(A)** ACC cell lines resemble CIMP-high ACC. Table includes list of recurrently altered genes identified in ACC-TCGA (Zheng *et al*., 2016). Third column describes the frequency of driver somatic alteration affecting that gene in CIMP-high ACC (Zheng *et al*., 2016). Genetic alterations for NCI-H295R and Y1 cell lines were identified and confirmed by inspecting BAM files from RNA-seq data generated by our group and others (Baba et al., 2014). Next-generation sequencing data is not available for ATC7L cell line and genetic alterations are therefore predicted based on known biological features of this line. Importantly, ATC7L is unlikely to harbor alterations leading to autonomous Wnt/β-catenin pathway activation since this line is very responsive to Wnt pathway stimulation (Walczak et al., 2014). **(B)** Heatmap of human adrenal microarray data from (Nishimoto *et al*., 2015). Genes selected based on the literature and involvement in pathways of interest. Genes encoding steroidogenic enzymes – *CYP11B1, HSD3B2*. Wnt/β-catenin pathway – *WNT4, AXIN2, LEF1, CTNNB1. G0S2* is targeted for methylation-dependent silencing in CIMP-high ACC, *NR5A1* encodes SF1, *MC2R* encodes the ACTH receptor, and *VSNL1* is a zG-specific gene upregulated in aldosterone-producing adrenocortical adenomas (Trejter et al., 2015; Williams et al., 2012). By this analysis, “zF genes”=*CYP11B1, HSD3B2, MC2R, G0S2* and “zG genes”=*VSNL1, WNT4, AXIN2, LEF1, CTNNB1*. **(C)** Y1 and ATC7L cells were fractionated into cytoplasmic (C) and nuclear (N) compartments, and protein lysates prepared from each compartment were analyzed by western blot, 15 μg protein per lane. For each epitope, Y1 and ATC7L lysates were run on the same gel/membrane. Irrelevant lanes between the sets of samples are cropped at the dashed line. Y1 express high levels of nuclear β-catenin, measured by total β-catenin in the top row, and active β-catenin (ABC) in second row; ATC7L express substantially lower levels of nuclear β-catenin. SF1 and PRC2 are nuclear and relatively comparable across both cell lines. H3K27ac and H3K27me3 are also comparable across cell lines and are shown here to demonstrate purity of cytoplasmic fraction. **(D)** ATC7L bears stronger zF differentiation than Y1. Top, heatmap of row z-score for qPCR data measuring expression of zonally expressed genes. Note that the genomic context of *Hsd3b1* (BLAST alignment with *HSD3B2* locus including upstream enhancer, data not shown) and its expression pattern render it the murine ortholog of *HSD3B2* (Simard et al., 2005). Bottom, average -ΔCt values for gene expression z-score calculated in top. Gene expression was measured by qPCR on reverse transcribed total mRNA from 3 independent biological replicates of ATC7L and Y1 cell lines grown in culture under standard conditions. **(E-F)** ATC7L (and not Y1) respond to Wnt pathway activation with partial zG differentiation at the expense of zF differentiation. Y1 (E, left) or ATC7L (E, right) were stimulated overnight (18.5-19.5 hrs) with LiCl and harvested for total mRNA and evaluation of gene expression by qPCR. LiCl is a potent and well characterized inhibitor of GSK3β, a core member of the destruction complex that targets β-catenin for degradation in the absence of Wnt ligands; LiCl administration mimics Wnt pathway activation (Stambolic et al., 1996). Concentration of LiCl in mM is indicated above each set of biological replicates (n=3 for Y1, n=2 for ATC7L) and heatmap is color coded by log2 of the fold change between the given concentration of LiCl over vehicle according to the legend below each plot. In E, right, ATC7L exhibits a dose dependent increase in expression of zG genes and canonical Wnt targets like *Axin2*, at the expense of zF genes. In contrast, Y1 (E, left) changes in gene expression in response to LiCl are minimal and encompass a broad upregulation of both zG and zF genes. In F, ATC7L and Y1 were stimulated with 20% Wnt3a conditioned medium (Wnt3a CM) or negative control (parental medium without Wnt3a, Par.), for 24 hours and harvested for total mRNA and evaluation of gene expression by qPCR. n=3 biological replicates for each cell line, and heatmap is color coded by log2 of the fold change between Wnt3a CM and Par. according to the legend below each plot. As in E, Wnt pathway activation could induce expression of zG genes in ATC7L only and had minimal effect on Y1 cells. **(G)** Y1 (and not ATC7L) exhibits exclusive induction of zF genes in response to PKA activation. Y1 and ATC7L were stimulated for 24-26 hours with 10 μM forskolin (FSK) or equivalent volume of vehicle (DMS) and harvested for total mRNA and evaluation of gene expression by qPCR. Forskolin is a well characterized stimulant of the PKA pathway, inducing intracellular accumulation of cAMP (Seamon *et al*., 1981). Heatmap is color coded by log2 of the fold change between FSK over DMS according to the legend below each plot, n=3 for each cell line. Treatment of adrenocortical cell lines with forskolin imitates the actions of ACTH (Xing *et al*., 2011). Forskolin administration in Y1 cells exclusively induced expression of zF genes; in contrast, forskolin administration in ATC7L cells induced expression of zF and zG genes. **(H)** Viability curves for Y1 and ATC7L treated with different classes of EZH2 inhibitors (EZH2i). Data are represented by mean (point) with SEM (whiskers). N=3 for each inhibitor per cell line. **(I)** Gene expression (fragments per kilobase of transcript per million mapped reads, FPKM) values from baseline NCI-H295R RNA-seq (n=3 replicates) of DNA methyltransferases (DNMT) *DNMT1, DNMT3A, DNMT3B*, and histone methyltransferases (HMT) *EZH1* and *EZH2*. **(J)** Example EZH2, H3K27me3, and H3K27ac ChIP-seq tracks across the *G0S2* locus in baseline NCI-H295R. Location of *G0S2* CpG island in black (top track), with region targeted for methylation in CIMP-high ACC depicted in grey (DMR, second track). Peak calls depicted by bars below each track.

**Supplementary Figure 4:**
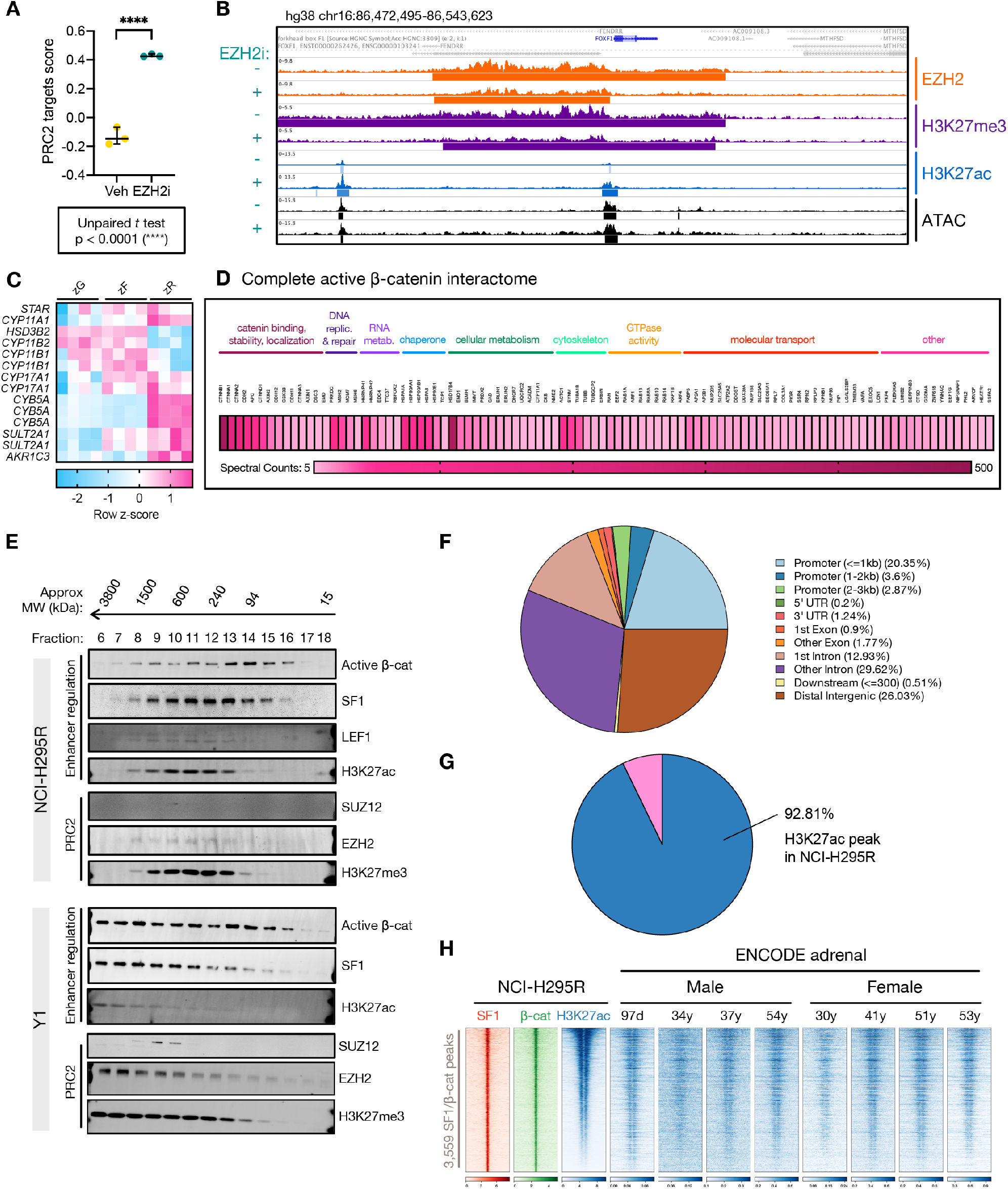
Related to Figures 3-4. **(A)** PRC2 targets score computed by GSVA (Hänzelmann *et al*., 2013) on RNA-seq data using BENPORATH_PRC2_TARGETS set (Ben-Porath *et al*., 2008) deposited in GSEA in baseline (Veh=vehicle-treated) or EZH2i-treated NCI-H295R. **(B)** Example EZH2, H3K27me3, H3K27ac, and ATAC-seq tracks across PRC2 target *FOXF1* locus in NCI-H295R at baseline (vehicle-treated, EZH2i-) or following EZH2 inhibition (EZH2i+). Peak calls are depicted by bars below each track. **(C)** Analysis of microarray data from (Nishimoto *et al*., 2015) demonstrates zonal expression of steroidogenic enzymes. **(D)** Peptides retrieved from active β-catenin-directed IP-MS on nuclear lysates from NCI-H295R. **(E)** Nuclear lysates from NCI-H295R or Y1 cell lines were fractionated by size exclusion chromatography. Equal volumes of each fraction in desired size range (15 kDa to 3800 kDa) were analyzed by western blot for epitopes shown right. Active β-cat=active β-catenin. As demonstrated in prior reports, PRC2 elution peaks at approximately 600 kDa (Margueron et al., 2008). N=2. **(F)** Characteristics of SF1/β-catenin binding sites in baseline NCI-H295R. 65% of peaks are >1000 bp away from a TSS and are therefore distal. **(G)** Pie chart depicting proportion of physiologic adrenal super-enhancers called by 3DIV analysis (Kim *et al*., 2020; Yang *et al*., 2018) on ENCODE samples (Consortium, 2012; Davis *et al*., 2018) overlapping with H3K27ac peaks in baseline NCI-H295R. **(H)** Heatmap of SF1, β-catenin, and H3K27ac signal in baseline NCI-H295R or H3K27ac signal across human adrenal samples deposited in ENCODE (Consortium, 2012; Davis *et al*., 2018) at 3,559 SF1/β-catenin cotargets, ranked by NCI-H295R H3K27ac signal. ENCODE samples from fetal male (97 day) and adult male and female (30-54 years of age). Centered at peak +/-3 kb window.

**Supplementary Figure 5:**
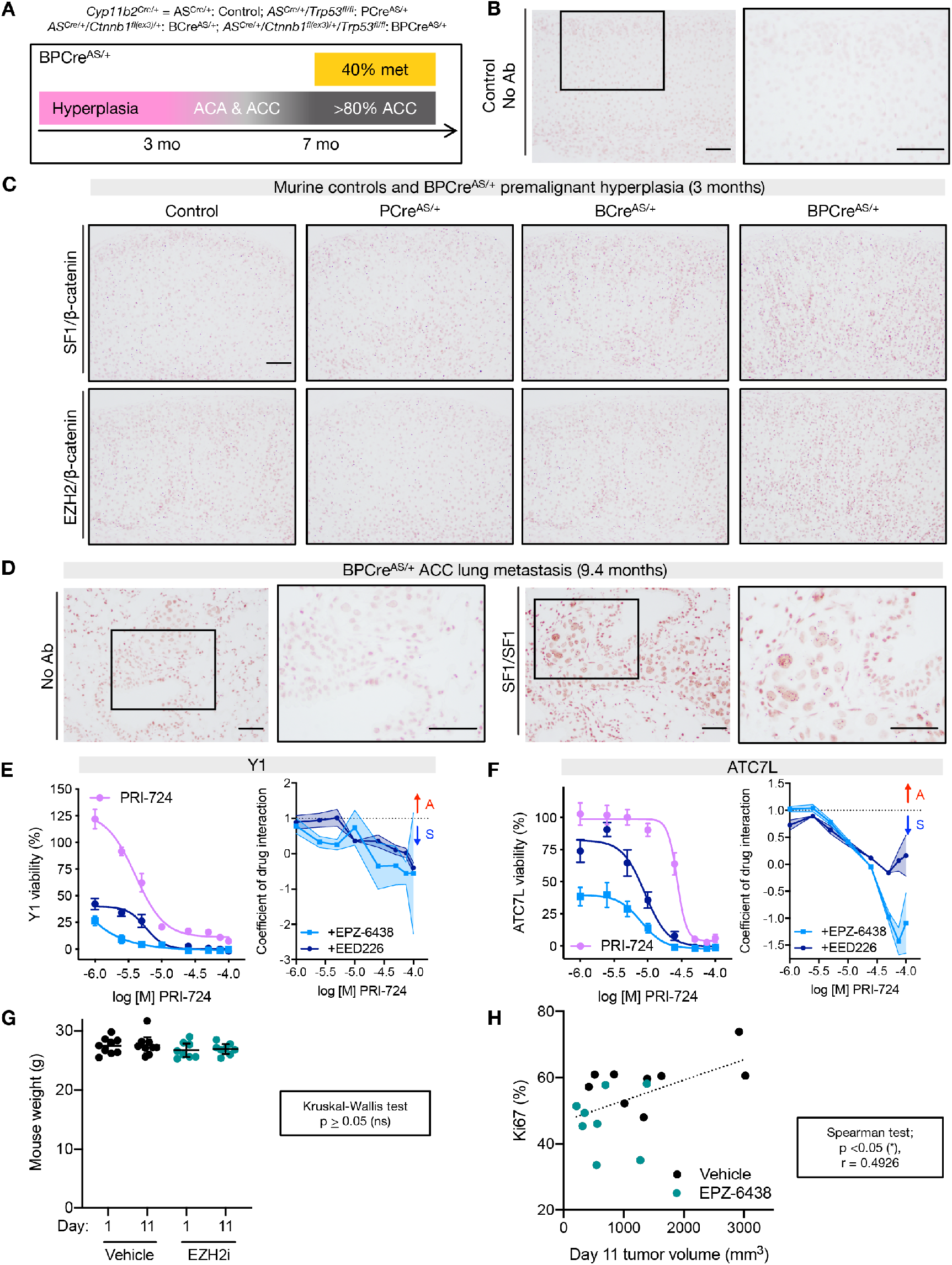
Related to Figures 5-7. **(A)** Top, key for abbreviations of mouse models used in the paper. Boxed, schematic of tumorigenesis in BPCre^AS/+^ model (Borges *et al*., 2020). By age 3 months, all mice exhibit adrenocortical hyperplasia. At age 5-7 months, approximately half of mice possess tumors (ACC and ACA). At 7-12 months, all mice possess tumors (with >80% meeting Weiss criteria for ACC) and 40% bear metastatic disease. **(B)** Example negative control PLA (no antibodies, No Ab) performed on sections from 3-month-old control (AS^Cre/+^) mice adrenals. Negative control PLA performed on n>10 murine adrenal and adrenal tumor tissue samples. Bar=100 μm. **(C)** Low magnification representative images of SF1/β-catenin and EZH2/β-catenin PLA performed on 3-month-old adrenals. Control, n=4 mice; PCre^AS/+^, n=3 mice; BCre^AS/+^, n=3 mice; BPCre^AS/+^, n=4 mice. Bar=100 μm. **(D)** Representative images of negative control (No Ab) and positive control (SF1/SF1) PLA performed on BPCre^AS/+^ lung metastases, n=3 mice. Bar=100 μm. **(E)** Left, viability curves for Y1 treated with increasing concentrations of CBP inhibitor (CBPi) PRI-724 +/-different classes of EZH2i (EPZ-6438 or EED226) at the IC-50 dose. Right, coefficient of drug interaction (CDI) for change in viability. CDI>1 represents antagonism (A), CDI <1 represents synergy (S). Data are represented by mean (point) with SEM (whiskers or error bands). CBPi, n=6; CBPi + EPZ-6438, n=3; CBPi + EED226, n=3. **(F)** Left, viability curves for ATC7L treated with increasing concentrations of CBPi PRI-724 +/-different classes of EZH2i at the IC-50 dose. Right, CDI for change in viability. Data are represented by mean (point) with SEM (whiskers or error bands). CBPi, n=3; CBPi + EPZ-6438, n=3; CBPi + EED226, n=3. **(G)** Mouse weight in BCH-ACC3A allograft experiment at beginning of experiment (Day 1) and end (Day 11) by treatment group. **(H)** Proliferation (measured by % nuclear Ki67 signal) vs final tumor volume across treatment groups in BCH-ACC3A allograft experiment.

## REFERENCES

Aryee, M.J., Jaffe, A.E., Corrada-Bravo, H., Ladd-Acosta, C., Feinberg, A.P., Hansen, K.D., and Irizarry, R.A. (2014). Minfi: a flexible and comprehensive Bioconductor package for the analysis of Infinium DNA methylation microarrays. Bioinformatics 30, 1363–1369. 10.1093/bioinformatics/btu049.

Assie, G., Letouze, E., Fassnacht, M., Jouinot, A., Luscap, W., Barreau, O., Omeiri, H., Rodriguez, S., Perlemoine, K., Rene-Corail, F., et al. (2014). Integrated genomic characterization of adrenocortical carcinoma. Nat Genet 46, 607–612. 10.1038/ng.2953.

Baba, T., Otake, H., Sato, T., Miyabayashi, K., Shishido, Y., Wang, C.Y., Shima, Y., Kimura, H., Yagi, M., Ishihara, Y., et al. (2014). Glycolytic genes are targets of the nuclear receptor Ad4BP/SF-1. Nat Commun 5, 3634. 10.1038/ncomms4634.

Basham, K.J., Rodriguez, S., Turcu, A.F., Lerario, A.M., Logan, C.Y., Rysztak, M.R., Gomez-Sanchez, C.E., Breault, D.T., Koo, B.K., Clevers, H., et al. (2019). A ZNRF3-dependent Wnt/β-catenin signaling gradient is required for adrenal homeostasis. Genes Dev 33, 209–220. 10.1101/gad.317412.118.

Bassett, M.H., White, P.C., and Rainey, W.E. (2004). A role for the NGFI-B family in adrenal zonation and adrenocortical disease. Endocr Res 30, 567–574. 10.1081/erc-200043715.

Baylin, S.B., and Jones, P.A. (2016). Epigenetic Determinants of Cancer. Cold Spring Harb Perspect Biol 8. 10.1101/cshperspect.a019505.

Bayliss, J., Mukherjee, P., Lu, C., Jain, S.U., Chung, C., Martinez, D., Sabari, B., Margol, A.S., Panwalkar, P., Parolia, A., et al. (2016). Lowered H3K27me3 and DNA hypomethylation define poorly prognostic pediatric posterior fossa ependymomas. Sci Transl Med 8, 366ra161. 10.1126/scitranslmed.aah6904.

Ben-Porath, I., Thomson, M.W., Carey, V.J., Ge, R., Bell, G.W., Regev, A., and Weinberg, R.A. (2008). An embryonic stem cell-like gene expression signature in poorly differentiated aggressive human tumors. Nat Genet 40, 499–507. 10.1038/ng.127.

Berest, I., Arnold, C., Reyes-Palomares, A., Palla, G., Rasmussen, K.D., Giles, H., Bruch, P.M., Huber, W., Dietrich, S., Helin, K., and Zaugg, J.B. (2019). Quantification of Differential Transcription Factor Activity and Multiomics-Based Classification into Activators and Repressors: diffTF. Cell Rep 29, 3147-3159.e3112. 10.1016/j.celrep.2019.10.106.

Biton, A., Bernard-Pierrot, I., Lou, Y., Krucker, C., Chapeaublanc, E., Rubio-Pérez, C., López-Bigas, N., Kamoun, A., Neuzillet, Y., Gestraud, P., et al. (2014). Independent component analysis uncovers the landscape of the bladder tumor transcriptome and reveals insights into luminal and basal subtypes. Cell Rep 9, 1235–1245. 10.1016/j.celrep.2014.10.035.

Borges, K.S., Pignatti, E., Leng, S., Kariyawasam, D., Ruiz-Babot, G., Ramalho, F.S., Taketo, M.M., Carlone, D.L., and Breault, D.T. (2020). Wnt/β-catenin activation cooperates with loss of p53 to cause adrenocortical carcinoma in mice. Oncogene 39, 5282–5291. 10.1038/s41388-020-1358-5.

Bracken, A.P., Pasini, D., Capra, M., Prosperini, E., Colli, E., and Helin, K. (2003). EZH2 is downstream of the pRB-E2F pathway, essential for proliferation and amplified in cancer. EMBO J 22, 5323–5335. 10.1093/emboj/cdg542.

Cao, J., Spielmann, M., Qiu, X., Huang, X., Ibrahim, D.M., Hill, A.J., Zhang, F., Mundlos, S., Christiansen, L., Steemers, F.J., et al. (2019). The single-cell transcriptional landscape of mammalian organogenesis. Nature 566, 496–502. 10.1038/s41586-019-0969-x.

Cao, R., and Zhang, Y. (2004). SUZ12 is required for both the histone methyltransferase activity and the silencing function of the EED-EZH2 complex. Mol Cell 15, 57–67. 10.1016/j.molcel.2004.06.020.

Catania, S., Dumesic, P.A., Pimentel, H., Nasif, A., Stoddard, C.I., Burke, J.E., Diedrich, J.K., Cook, S., Shea, T., Geinger, E., et al. (2020). Evolutionary Persistence of DNA Methylation for Millions of Years after Ancient Loss of a De Novo Methyltransferase. Cell 180, 263-277.e220. 10.1016/j.cell.2019.12.012.

Cerami, E., Gao, J., Dogrusoz, U., Gross, B.E., Sumer, S.O., Aksoy, B.A., Jacobsen, A., Byrne, C.J., Heuer, M.L., Larsson, E., et al. (2012). The cBio cancer genomics portal: an open platform for exploring multi-dimensional cancer genomics data. Cancer Discov 2, 401–404. 10.1158/2159-8290.CD-12-0095.

Chaligne, R., Gaiti, F., Silverbush, D., Schiffman, J.S., Weisman, H.R., Kluegel, L., Gritsch, S., Deochand, S.D., Gonzalez Castro, L.N., Richman, A.R., et al. (2021). Epigenetic encoding, heritability and plasticity of glioma transcriptional cell states. Nat Genet 53, 1469–1479. 10.1038/s41588-021-00927-7.

Chung, C., Sweha, S.R., Pratt, D., Tamrazi, B., Panwalkar, P., Banda, A., Bayliss, J., Hawes, D., Yang, F., Lee, H.J., et al. (2020). Integrated Metabolic and Epigenomic Reprograming by H3K27M Mutations in Diffuse Intrinsic Pontine Gliomas. Cancer Cell. 10.1016/j.ccell.2020.07.008.

Consortium, E.P. (2012). An integrated encyclopedia of DNA elements in the human genome. Nature 489, 57–74. 10.1038/nature11247.

Corces, M.R., Granja, J.M., Shams, S., Louie, B.H., Seoane, J.A., Zhou, W., Silva, T.C., Groeneveld, C., Wong, C.K., Cho, S.W., et al. (2018). The chromatin accessibility landscape of primary human cancers. Science 362. 10.1126/science.aav1898.

Corces, M.R., Trevino, A.E., Hamilton, E.G., Greenside, P.G., Sinnott-Armstrong, N.A., Vesuna, S., Satpathy, A.T., Rubin, A.J., Montine, K.S., Wu, B., et al. (2017). An improved ATAC-seq protocol reduces background and enables interrogation of frozen tissues. Nature Methods 14, 959–962.

Davis, C.A., Hitz, B.C., Sloan, C.A., Chan, E.T., Davidson, J.M., Gabdank, I., Hilton, J.A., Jain, K., Baymuradov, U.K., Narayanan, A.K., et al. (2018). The Encyclopedia of DNA elements (ENCODE): data portal update. Nucleic Acids Res 46, D794–D801. 10.1093/nar/gkx1081.

Deevy, O., and Bracken, A.P. (2019). PRC2 functions in development and congenital disorders. Development 146. 10.1242/dev.181354.

DeLuca, D.S., Levin, J.Z., Sivachenko, A., Fennell, T., Nazaire, M.D., Williams, C., Reich, M., Winckler, W., and Getz, G. (2012). RNA-SeQC: RNA-seq metrics for quality control and process optimization. Bioinformatics 28, 1530–1532. 10.1093/bioinformatics/bts196.

Dobin, A., Davis, C.A., Schlesinger, F., Drenkow, J., Zaleski, C., Jha, S., Batut, P., Chaisson, M., and Gingeras, T.R. (2013). STAR: ultrafast universal RNA-seq aligner. Bioinformatics 29, 15–21. 10.1093/bio-informatics/bts635.

Domcke, S., Hill, A.J., Daza, R.M., Cao, J., O’Day, D.R., Pliner, H.A., Al-dinger, K.A., Pokholok, D., Zhang, F., Milbank, J.H., et al. (2020). A human cell atlas of fetal chromatin accessibility. Science 370. 10.1126/science.aba7612.

Drelon, C., Berthon, A., Mathieu, M., Ragazzon, B., Kuick, R., Tabbal, H., Septier, A., Rodriguez, S., Batisse-Lignier, M., Sahut-Barnola, I., et al. (2016). EZH2 is overexpressed in adrenocortical carcinoma and is associated with disease progression. Hum Mol Genet 25, 2789–2800. 10.1093/hmg/ddw136.

Easwaran, H., Johnstone, S.E., Van Neste, L., Ohm, J., Mosbruger, T., Wang, Q., Aryee, M.J., Joyce, P., Ahuja, N., Weisenberger, D., et al. (2012). A DNA hypermethylation module for the stem/progenitor cell signature of cancer. Genome Res 22, 837–849. 10.1101/gr.131169.111.

Finco, I., Lerario, A.M., and Hammer, G.D. (2018). Sonic Hedgehog and WNT Signaling Promote Adrenal Gland Regeneration in Male Mice. Endocrinology 159, 579–596. 10.1210/en.2017-03061.

Flavahan, W.A., Gaskell, E., and Bernstein, B.E. (2017). Epigenetic plasticity and the hallmarks of cancer. Science 357. 10.1126/sci-ence.aal2380.

Gal-Yam, E.N., Egger, G., Iniguez, L., Holster, H., Einarsson, S., Zhang, X., Lin, J.C., Liang, G., Jones, P.A., and Tanay, A. (2008). Frequent switching of Polycomb repressive marks and DNA hypermethylation in the PC3 prostate cancer cell line. Proc Natl Acad Sci U S A 105, 12979–12984. 10.1073/pnas.0806437105.

Gao, J., Aksoy, B.A., Dogrusoz, U., Dresdner, G., Gross, B., Sumer, S.O., Sun, Y., Jacobsen, A., Sinha, R., Larsson, E., et al. (2013). Integrative analysis of complex cancer genomics and clinical profiles using the cBioPortal. Sci Signal 6, pl1. 10.1126/scisignal.2004088.

Gaspar, J.M. (2018). Genrich: detecting sites of genomic enrichment. https://github.com/jsh58/Genrich.

Gummow, B.M., Winnay, J.N., and Hammer, G.D. (2003). Convergence of Wnt signaling and steroidogenic factor-1 (SF-1) on transcription of the rat inhibin alpha gene. J Biol Chem 278, 26572–26579. 10.1074/jbc.M212677200.

Han, X., Zhou, Z., Fei, L., Sun, H., Wang, R., Chen, Y., Chen, H., Wang, J., Tang, H., Ge, W., et al. (2020). Construction of a human cell landscape at single-cell level. Nature 581, 303–309. 10.1038/s41586-020-2157-4.

Heinz, S., Benner, C., Spann, N., Bertolino, E., Lin, Y.C., Laslo, P., Cheng, J.X., Murre, C., Singh, H., and Glass, C.K. (2010). Simple combinations of lineage-determining transcription factors prime cis-regulatory elements required for macrophage and B cell identities. Mol Cell 38, 576–589. 10.1016/j.molcel.2010.05.004.

Hnisz, D., Abraham, B.J., Lee, T.I., Lau, A., Saint-André, V., Sigova, A.A., Hoke, H.A., and Young, R.A. (2013). Super-enhancers in the control of cell identity and disease. Cell 155, 934–947. 10.1016/j.cell.2013.09.053.

Hnisz, D., Schuijers, J., Lin, C.Y., Weintraub, A.S., Abraham, B.J., Lee, T.I., Bradner, J.E., and Young, R.A. (2015). Convergence of developmental and oncogenic signaling pathways at transcriptional super-enhancers. Mol Cell 58, 362–370. 10.1016/j.mol-cel.2015.02.014.

Hoffmeyer, K., Junghans, D., Kanzler, B., and Kemler, R. (2017). Trimethylation and Acetylation of β-Catenin at Lysine 49 Represent Key Elements in ESC Pluripotency. Cell Rep 18, 2815–2824. 10.1016/j.celrep.2017.02.076.

Hossain, A., and Saunders, G.F. (2003). Synergistic cooperation between the beta-catenin signaling pathway and steroidogenic factor 1 in the activation of the Mullerian inhibiting substance type II receptor. J Biol Chem 278, 26511–26516. 10.1074/jbc.M300804200.

Hovestadt, V., and Zapatka, M. (2020). conumee: Enhanced copy-number variation analysis using Illumina DNA methylation arrays.

Hänzelmann, S., Castelo, R., and Guinney, J. (2013). GSVA: gene set variation analysis for microarray and RNA-seq data. BMC Bioinformatics 14, 7. 10.1186/1471-2105-14-7.

Ireland, A.S., Micinski, A.M., Kastner, D.W., Guo, B., Wait, S.J., Spainhower, K.B., Conley, C.C., Chen, O.S., Guthrie, M.R., Soltero, D., et al. (2020). MYC Drives Temporal Evolution of Small Cell Lung Cancer Subtypes by Reprogramming Neuroendocrine Fate. Cancer Cell 38, 60-78.e12. 10.1016/j.ccell.2020.05.001.

Irizarry, R.A., Hobbs, B., Collin, F., Beazer-Barclay, Y.D., Antonellis, K.J., Scherf, U., and Speed, T.P. (2003). Exploration, normalization, and summaries of high density oligonucleotide array probe level data. Biostatistics 4, 249–264. 10.1093/biostatistics/4.2.249.

Jung, I., Schmitt, A., Diao, Y., Lee, A.J., Liu, T., Yang, D., Tan, C., Eom, J., Chan, M., Chee, S., et al. (2019). A compendium of promotercentered long-range chromatin interactions in the human genome. Nat Genet 51, 1442–1449. 10.1038/s41588-019-0494-8.

Kadoch, C., Hargreaves, D.C., Hodges, C., Elias, L., Ho, L., Ranish, J., and Crabtree, G.R. (2013). Proteomic and bioinformatic analysis of mammalian SWI/SNF complexes identifies extensive roles in human malignancy. Nat Genet 45, 592–601. 10.1038/ng.2628.

Kahn, M. (2014). Can we safely target the WNT pathway? Nat Rev Drug Discov 13, 513–532. 10.1038/nrd4233.

Kennell, J.A., O’Leary, E.E., Gummow, B.M., Hammer, G.D., and Mac-Dougald, O.A. (2003). T-cell factor 4N (TCF-4N), a novel isoform of mouse TCF-4, synergizes with beta-catenin to coactivate C/EBPalpha and steroidogenic factor 1 transcription factors. Mol Cell Biol 23, 5366–5375. 10.1128/MCB.23.15.5366-5375.2003.

Kim, K., Jang, I., Kim, M., Choi, J., Kim, M.S., Lee, B., and Jung, I. (2020). 3DIV update for 2021: a comprehensive resource of 3D genome and 3D cancer genome. Nucleic Acids Res. 10.1093/nar/gkaa1078.

LaFave, L.M., Kartha, V.K., Ma, S., Meli, K., Del Priore, I., Lareau, C., Naranjo, S., Westcott, P.M.K., Duarte, F.M., Sankar, V., et al. (2020). Epigenomic State Transitions Characterize Tumor Progression in Mouse Lung Adenocarcinoma. Cancer Cell 38, 212-228.e213. 10.1016/j.ccell.2020.06.006.

Langmead, B., and Salzberg, S.L. (2012). Fast gapped-read alignment with Bowtie 2. Nat Methods 9, 357–359. 10.1038/nmeth.1923.

Lawrence, M., Gentleman, R., and Carey, V. (2009). rtracklayer: an R package for interfacing with genome browsers. Bioinformatics 25, 1841–1842. 10.1093/bioinformatics/btp328.

Lerario, A.M., Mohan, D.R., and Hammer, G.D. (2022). Update on Biology and Genomics of Adrenocortical Carcinomas: Rationale for Emerging Therapies. Endocrine Reviews. 10.1210/endrev/bnac012.

Li, H., Liefke, R., Jiang, J., Kurland, J.V., Tian, W., Deng, P., Zhang, W., He, Q., Patel, D.J., Bulyk, M.L., et al. (2017). Polycomb-like proteins link the PRC2 complex to CpG islands. Nature 549, 287–291. 10.1038/nature23881.

Liao, Y., Smyth, G.K., and Shi, W. (2014). featureCounts: an efficient general purpose program for assigning sequence reads to genomic features. Bioinformatics 30, 923–930. 10.1093/bioinformat-ics/btt656.

Liberzon, A., Subramanian, A., Pinchback, R., Thorvaldsdóttir, H., Tamayo, P., and Mesirov, J.P. (2011). Molecular signatures database (MSigDB) 3.0. Bioinformatics 27, 1739–1740. 10.1093/bioinfor-matics/btr260.

Lovén, J., Hoke, H.A., Lin, C.Y., Lau, A., Orlando, D.A., Vakoc, C.R., Bradner, J.E., Lee, T.I., and Young, R.A. (2013). Selective inhibition of tumor oncogenes by disruption of super-enhancers. Cell 153, 320–334. 10.1016/j.cell.2013.03.036.

Margueron, R., Justin, N., Ohno, K., Sharpe, M.L., Son, J., Drury, W.J., Voigt, P., Martin, S.R., Taylor, W.R., De Marco, V., et al. (2009). Role of the polycomb protein EED in the propagation of repressive histone marks. Nature 461, 762–767. 10.1038/nature08398.

Margueron, R., Li, G., Sarma, K., Blais, A., Zavadil, J., Woodcock, C.L., Dynlacht, B.D., and Reinberg, D. (2008). Ezh1 and Ezh2 maintain repressive chromatin through different mechanisms. Mol Cell 32, 503–518. 10.1016/j.molcel.2008.11.004.

Mathieu, M., Drelon, C., Rodriguez, S., Tabbal, H., Septier, A., Damon-Soubeyrand, C., Dumontet, T., Berthon, A., Sahut-Barnola, I., Djari, C., et al. (2018). Steroidogenic differentiation and PKA signaling are programmed by histone methyltransferase EZH2 in the adrenal cortex. Proc Natl Acad Sci U S A 115, E12265–E12274. 10.1073/pnas.1809185115.

McCarthy, D.J., Chen, Y., and Smyth, G.K. (2012). Differential expression analysis of multifactor RNA-Seq experiments with respect to biological variation. Nucleic Acids Res 40, 4288–4297. 10.1093/nar/gks042.

Merika, M., Williams, A.J., Chen, G., Collins, T., and Thanos, D. (1998). Recruitment of CBP/p300 by the IFN beta enhanceosome is required for synergistic activation of transcription. Mol Cell 1, 277–287. 10.1016/s1097-2765(00)80028-3.

Mizusaki, H., Kawabe, K., Mukai, T., Ariyoshi, E., Kasahara, M., Yoshioka, H., Swain, A., and Morohashi, K. (2003). Dax-1 (dosage-sensitive sex reversal-adrenal hypoplasia congenita critical region on the X chromosome, gene 1) gene transcription is regulated by wnt4 in the female developing gonad. Mol Endocrinol 17, 507–519. 10.1210/me.2002-0362.

Mohan, D.R., Lerario, A.M., Else, T., Mukherjee, B., Almeida, M.Q., Vinco, M., Rege, J., Mariani, B.M.P., Zerbini, M.C.N., Mendonca, B.B., et al. (2019). Targeted Assessment of G0S2 Methylation Identifies a Rapidly Recurrent, Routinely Fatal Molecular Subtype of Adrenocortical Carcinoma. Clin Cancer Res 25, 3276–3288. 10.1158/1078-0432.CCR-18-2693.

Montgomery, N.D., Yee, D., Chen, A., Kalantry, S., Chamberlain, S.J., Otte, A.P., and Magnuson, T. (2005). The murine polycomb group protein Eed is required for global histone H3 lysine-27 methylation. Curr Biol 15, 942–947. 10.1016/j.cub.2005.04.051.

Mootha, V.K., Lindgren, C.M., Eriksson, K.F., Subramanian, A., Sihag, S., Lehar, J., Puigserver, P., Carlsson, E., Ridderstråle, M., Laurila, E., et al. (2003). PGC-1alpha-responsive genes involved in oxidative phosphorylation are coordinately downregulated in human diabetes. Nat Genet 34, 267–273. 10.1038/ng1180.

Morschhauser, F., Tilly, H., Chaidos, A., McKay, P., Phillips, T., Assouline, S., Batlevi, C.L., Campbell, P., Ribrag, V., Damaj, G.L., et al. (2020). Tazemetostat for patients with relapsed or refractory follicular lymphoma: an open-label, single-arm, multicentre, phase 2 trial. Lancet Oncol 21, 1433–1442. 10.1016/S1470-2045(20)30441-1.

NCI (2005-2018). The Cancer Genome Atlas. https://www.can-cer.gov/tcga.

Neri, F., Krepelova, A., Incarnato, D., Maldotti, M., Parlato, C., Galvagni, F., Matarese, F., Stunnenberg, H.G., and Oliviero, S. (2013). Dnmt3L antagonizes DNA methylation at bivalent promoters and favors DNA methylation at gene bodies in ESCs. Cell 155, 121–134. 10.1016/j.cell.2013.08.056.

Nishimoto, K., Tomlins, S.A., Kuick, R., Cani, A.K., Giordano, T.J., Hovelson, D.H., Liu, C.J., Sanjanwala, A.R., Edwards, M.A., Gomez-Sanchez, C.E., et al. (2015). Aldosterone-stimulating somatic gene mutations are common in normal adrenal glands. Proc Natl Acad Sci U S A 112, E4591–4599. 10.1073/pnas.1505529112.

Nusse, R., and Clevers, H. (2017). Wnt/β-Catenin Signaling, Disease, and Emerging Therapeutic Modalities. Cell 169, 985–999. 10.1016/j.cell.2017.05.016.

Ohm, J.E., McGarvey, K.M., Yu, X., Cheng, L., Schuebel, K.E., Cope, L., Mohammad, H.P., Chen, W., Daniel, V.C., Yu, W., et al. (2007). A stem cell-like chromatin pattern may predispose tumor suppressor genes to DNA hypermethylation and heritable silencing. Nat Genet 39, 237–242. 10.1038/ng1972.

Panwalkar, P., Clark, J., Ramaswamy, V., Hawes, D., Yang, F., Dunham, C., Yip, S., Hukin, J., Sun, Y., Schipper, M.J., et al. (2017). Immuno-histochemical analysis of H3K27me3 demonstrates global reduction in group-A childhood posterior fossa ependymoma and is a powerful predictor of outcome. Acta Neuropathol. 10.1007/s00401-017-1752-4.

Pasini, D., Bracken, A.P., Jensen, M.R., Lazzerini Denchi, E., and Helin, K. (2004). Suz12 is essential for mouse development and for EZH2 histone methyltransferase activity. EMBO J 23, 4061–4071. 10.1038/sj.emboj.7600402.

Perino, M., van Mierlo, G., Karemaker, I.D., van Genesen, S., Vermeulen, M., Marks, H., van Heeringen, S.J., and Veenstra, G.J.C. (2018). MTF2 recruits Polycomb Repressive Complex 2 by helical-shape-selective DNA binding. Nat Genet 50, 1002–1010. 10.1038/s41588-018-0134-8.

Pham, T.N.D., Kumar, K., DeCant, B.T., Shang, M., Munshi, S.Z., Matsangou, M., Ebine, K., and Munshi, H.G. (2019). Induction of MNK Kinase-dependent eIF4E Phosphorylation by Inhibitors Targeting BET Proteins Limits Efficacy of BET Inhibitors. Mol Cancer Ther 18, 235–244. 10.1158/1535-7163.MCT-18-0768.

Pignatti, E., Leng, S., Yuchi, Y., Borges, K.S., Guagliardo, N.A., Shah, M.S., Ruiz-Babot, G., Kariyawasam, D., Taketo, M.M., Miao, J., et al. (2020). Beta-Catenin Causes Adrenal Hyperplasia by Blocking Zonal Transdifferentiation. Cell Rep 31, 107524. 10.1016/j.celrep.2020.107524.

Pott, S., and Lieb, J.D. (2015). What are super-enhancers? Nat Genet 47, 8–12. 10.1038/ng.3167.

Qiu, X., Hill, A., Packer, J., Lin, D., Ma, Y.A., and Trapnell, C. (2017a). Single-cell mRNA quantification and differential analysis with Census. Nat Methods 14, 309–315. 10.1038/nmeth.4150.

Qiu, X., Mao, Q., Tang, Y., Wang, L., Chawla, R., Pliner, H.A., and Trapnell, C. (2017b). Reversed graph embedding resolves complex single-cell trajectories. Nat Methods 14, 979–982. 10.1038/nmeth.4402.

Quinlan, A.R., and Hall, I.M. (2010). BEDTools: a flexible suite of utilities for comparing genomic features. Bioinformatics 26, 841–842. 10.1093/bioinformatics/btq033.

Raisner, R., Kharbanda, S., Jin, L., Jeng, E., Chan, E., Merchant, M., Haverty, P.M., Bainer, R., Cheung, T., Arnott, D., et al. (2018). Enhancer Activity Requires CBP/P300 Bromodomain-Dependent Histone H3K27 Acetylation. Cell Rep 24, 1722–1729. 10.1016/j.celrep.2018.07.041.

Ramírez, F., Ryan, D.P., Grüning, B., Bhardwaj, V., Kilpert, F., Richter, A.S., Heyne, S., Dündar, F., and Manke, T. (2016). deepTools2: a next generation web server for deep-sequencing data analysis. Nucleic Acids Res 44, W160–165. 10.1093/nar/gkw257.

Ritchie, M.E., Phipson, B., Wu, D., Hu, Y., Law, C.W., Shi, W., and Smyth, G.K. (2015). limma powers differential expression analyses for RNA-sequencing and microarray studies. Nucleic Acids Res 43, e47. 10.1093/nar/gkv007.

Robinson, M.D., McCarthy, D.J., and Smyth, G.K. (2010). edgeR: a Bioconductor package for differential expression analysis of digital gene expression data. Bioinformatics 26, 139–140. 10.1093/bioin-formatics/btp616.

Ross-Innes, C.S., Stark, R., Teschendorff, A.E., Holmes, K.A., Ali, H.R., Dunning, M.J., Brown, G.D., Gojis, O., Ellis, I.O., Green, A.R., et al. (2012). Differential oestrogen receptor binding is associated with clinical outcome in breast cancer. Nature 481, 389–393. 10.1038/nature10730.

Sabari, B.R., Dall’Agnese, A., Boija, A., Klein, I.A., Coffey, E.L., Shrinivas, K., Abraham, B.J., Hannett, N.M., Zamudio, A.V., Manteiga, J.C., et al. (2018). Coactivator condensation at super-enhancers links phase separation and gene control. Science 361. 10.1126/sci-ence.aar3958.

Schindelin, J., Arganda-Carreras, I., Frise, E., Kaynig, V., Longair, M., Pietzsch, T., Preibisch, S., Rueden, C., Saalfeld, S., Schmid, B., et al. (2012). Fiji: an open-source platform for biological-image analysis. Nat Methods 9, 676–682. 10.1038/nmeth.2019.

Schneider, C.A., Rasband, W.S., and Eliceiri, K.W. (2012). NIH Image to ImageJ: 25 years of image analysis. Nat Methods 9, 671–675. 10.1038/nmeth.2089.

Schuettengruber, B., Bourbon, H.M., Di Croce, L., and Cavalli, G. (2017). Genome Regulation by Polycomb and Trithorax: 70 Years and Counting. Cell 171, 34–57. 10.1016/j.cell.2017.08.002.

Schwitalla, S., Fingerle, A.A., Cammareri, P., Nebelsiek, T., Göktuna, S.I., Ziegler, P.K., Canli, O., Heijmans, J., Huels, D.J., Moreaux, G., et al. (2013). Intestinal tumorigenesis initiated by dedifferentiation and acquisition of stem-cell-like properties. Cell 152, 25–38. 10.1016/j.cell.2012.12.012.

Seamon, K.B., Padgett, W., and Daly, J.W. (1981). Forskolin: unique diterpene activator of adenylate cyclase in membranes and in intact cells. Proc Natl Acad Sci U S A 78, 3363–3367. 10.1073/pnas.78.6.3363.

Shibamoto, S., Higano, K., Takada, R., Ito, F., Takeichi, M., and Takada, S. (1998). Cytoskeletal reorganization by soluble Wnt-3a protein signalling. Genes Cells 3, 659–670. 10.1046/j.13652443.1998.00221.x.

Shpynov, O., Dievskii, A., Chernyatchik, R., Tsurinov, P., and Artyomov, M.N. (2021). Semi-supervised peak calling with SPAN and JBR Genome Browser. Bioinformatics. 10.1093/bioinformatics/btab376.

Simard, J., Ricketts, M.L., Gingras, S., Soucy, P., Feltus, F.A., and Melner, M.H. (2005). Molecular biology of the 3beta-hydroxysteroid dehy-drogenase/delta5-delta4 isomerase gene family. Endocr Rev 26, 525–582. 10.1210/er.2002-0050.

Stambolic, V., Ruel, L., and Woodgett, J.R. (1996). Lithium inhibits gly-cogen synthase kinase-3 activity and mimics wingless signalling in intact cells. Curr Biol 6, 1664–1668. 10.1016/s0960-9822(02)70790-2.

Stark, R., and Brown, G. (2011). DiffBind: differential binding analysis of ChIP-Seq peak data. 10.18129/B9.bioc.DiffBind.

Stuart, T., Butler, A., Hoffman, P., Hafemeister, C., Papalexi, E., Mauck, W.M., Hao, Y., Stoeckius, M., Smibert, P., and Satija, R. (2019). Comprehensive Integration of Single-Cell Data. Cell 177, 1888-1902.e1821. 10.1016/j.cell.2019.05.031.

Stuart, T., Srivastava, A., Madad, S., Lareau, C.A., and Satija, R. (2021). Single-cell chromatin state analysis with Signac. Nat Methods 18, 1333–1341. 10.1038/s41592-021-01282-5.

Subramanian, A., Tamayo, P., Mootha, V.K., Mukherjee, S., Ebert, B.L., Gillette, M.A., Paulovich, A., Pomeroy, S.L., Golub, T.R., Lander, E.S., and Mesirov, J.P. (2005). Gene set enrichment analysis: a knowledge-based approach for interpreting genome-wide expres-sion profiles. Proc Natl Acad Sci U S A 102, 15545–15550. 10.1073/pnas.0506580102.

Tang, Z., Li, C., Kang, B., Gao, G., and Zhang, Z. (2017). GEPIA: a web server for cancer and normal gene expression profiling and interactive analyses. Nucleic Acids Res 45, W98–W102. 10.1093/nar/gkx247.

Tao, Y., Kang, B., Petkovich, D.A., Bhandari, Y.R., In, J., Stein-O’Brien, G., Kong, X., Xie, W., Zachos, N., Maegawa, S., et al. (2019). Aginglike Spontaneous Epigenetic Silencing Facilitates Wnt Activation, Stemness, and Braf. Cancer Cell 35, 315-328.e316. 10.1016/j.ccell.2019.01.005.

Team, R.C. (2016). R: A language and environment for statistical computing. https://www.r-project.org/.

Toyota, M., Ahuja, N., Ohe-Toyota, M., Herman, J.G., Baylin, S.B., and Issa, J.P. (1999). CpG island methylator phenotype in colorectal cancer. Proc Natl Acad Sci U S A 96, 8681–8686. 10.1073/pnas.96.15.8681.

Trapnell, C., Cacchiarelli, D., Grimsby, J., Pokharel, P., Li, S., Morse, M., Lennon, N.J., Livak, K.J., Mikkelsen, T.S., and Rinn, J.L. (2014). The dynamics and regulators of cell fate decisions are revealed by pseudotemporal ordering of single cells. Nat Biotechnol 32, 381–386. 10.1038/nbt.2859.

Trejter, M., Hochol, A., Tyczewska, M., Ziolkowska, A., Jopek, K., Szyszka, M., Malendowicz, L.K., and Rucinski, M. (2015). Visinin-like peptide 1 in adrenal gland of the rat. Gene expression and its hormonal control. Peptides 63, 22–29. 10.1016/j.pep-tides.2014.10.017.

van Iterson, M., Tobi, E.W., Slieker, R.C., den Hollander, W., Luijk, R., Slagboom, P.E., and Heijmans, B.T. (2014). MethylAid: visual and interactive quality control of large Illumina 450k datasets. Bioinformatics 30, 3435–3437. 10.1093/bioinformatics/btu566.

Vaz, M., Hwang, S.Y., Kagiampakis, I., Phallen, J., Patil, A., O’Hagan, H.M., Murphy, L., Zahnow, C.A., Gabrielson, E., Velculescu, V.E., et al. (2017). Chronic Cigarette Smoke-Induced Epigenomic Changes Precede Sensitization of Bronchial Epithelial Cells to Single-Step Transformation by KRAS Mutations. Cancer Cell 32, 360-376.e366. 10.1016/j.ccell.2017.08.006.

Viré, E., Brenner, C., Deplus, R., Blanchon, L., Fraga, M., Didelot, C., Morey, L., Van Eynde, A., Bernard, D., Vanderwinden, J.M., et al. (2006). The Polycomb group protein EZH2 directly controls DNA methylation. Nature 439, 871–874. 10.1038/nature04431.

Walczak, E.M., Kuick, R., Finco, I., Bohin, N., Hrycaj, S.M., Wellik, D.M., and Hammer, G.D. (2014). Wnt signaling inhibits adrenal steroido-genesis by cell-autonomous and non-cell-autonomous mechanisms. Mol Endocrinol 28, 1471–1486. 10.1210/me.2014-1060.

Warde, K.M., Liu, L., Smith, L.J., Lohman, B.K., Stubben, C.J., Ekiz, H.A., Ammer, J.L., Converso-Baran, K., Giordano, T.J., Hammer, G.D., and Basham, K.J. (2022). Senescence-Induced Immune Remodeling Facilitates Metastatic Adrenal Cancer in a Sex-Dimorphic Manner. bioRxiv, 2022.2004.2029.488426. 10.1101/2022.04.29.488426.

Whyte, W.A., Orlando, D.A., Hnisz, D., Abraham, B.J., Lin, C.Y., Kagey, M.H., Rahl, P.B., Lee, T.I., and Young, R.A. (2013). Master tran-scription factors and mediator establish super-enhancers at key cell identity genes. Cell 153, 307–319. 10.1016/j.cell.2013.03.035.

Widschwendter, M., Fiegl, H., Egle, D., Mueller-Holzner, E., Spizzo, G., Marth, C., Weisenberger, D.J., Campan, M., Young, J., Jacobs, I., and Laird, P.W. (2007). Epigenetic stem cell signature in cancer. Nat Genet 39, 157–158. 10.1038/ng1941.

Williams, T.A., Monticone, S., Crudo, V., Warth, R., Veglio, F., and Mu-latero, P. (2012). Visinin-like 1 is upregulated in aldosterone-pro-ducing adenomas with KCNJ5 mutations and protects from calcium-induced apoptosis. Hypertension 59, 833–839. 10.1161/HYPER-TENSIONAHA.111.188532.

Wilmouth, J.J., Olabe, J., Garcia-Garcia, D., Lucas, C., Guiton, R., Roucher-Boulez, F., Dufour, D., Damon-Soubeyrand, C., Sahut-Barnola, I., Pointud, J.-C., et al. (2022). Sexually dimorphic activation of innate antitumour immunity prevents adrenocortical carcinoma development. bioRxiv, 2022.2004.2029.489846. 10.1101/2022.04.29.489846.

Wright, C., O’Day, P., Alyamani, M., Sharifi, N., and Auchus, R.J. (2020). Abiraterone acetate treatment lowers 11-oxygenated androgens. Eur J Endocrinol 182, 413–421. 10.1530/EJE-19-0905.

Xing, Y., Edwards, M.A., Ahlem, C., Kennedy, M., Cohen, A., Gomez-Sanchez, C.E., and Rainey, W.E. (2011). The effects of ACTH on steroid metabolomic profiles in human adrenal cells. J Endocrinol 209, 327–335. 10.1530/JOE-10-0493.

Yakulov, T., Raggioli, A., Franz, H., and Kemler, R. (2013). Wnt3a-dependent and -independent protein interaction networks of chromatin-bound β-catenin in mouse embryonic stem cells. Mol Cell Proteomics 12, 1980–1994. 10.1074/mcp.M112.026914.

Yang, D., Jang, I., Choi, J., Kim, M.S., Lee, A.J., Kim, H., Eom, J., Kim, D., Jung, I., and Lee, B. (2018). 3DIV: A 3D-genome Interaction Viewer and database. Nucleic Acids Res 46, D52–D57. 10.1093/nar/gkx1017.

Yu, G., Wang, L.G., and He, Q.Y. (2015). ChIPseeker: an R/Bioconductor package for ChIP peak annotation, comparison and visualization. Bioinformatics 31, 2382–2383. 10.1093/bioinformatics/btv145.

Zamudio, A.V., Dall’Agnese, A., Henninger, J.E., Manteiga, J.C., Afeyan, L.K., Hannett, N.M., Coffey, E.L., Li, C.H., Oksuz, O., Sabari, B.R., et al. (2019). Mediator Condensates Localize Signaling Factors to Key Cell Identity Genes. Mol Cell 76, 753-766.e756. 10.1016/j.molcel.2019.08.016.

Zhang, K., Hocker, J.D., Miller, M., Hou, X., Chiou, J., Poirion, O.B., Qiu, Y., Li, Y.E., Gaulton, K.J., Wang, A., et al. (2021). A single-cell atlas of chromatin accessibility in the human genome. Cell 184, 5985-6001.e5919. 10.1016/j.cell.2021.10.024.

Zheng, S., Cherniack, A.D., Dewal, N., Moffitt, R.A., Danilova, L., Murray, B.A., Lerario, A.M., Else, T., Knijnenburg, T.A., Ciriello, G., et al. (2016). Comprehensive Pan-Genomic Characterization of Adreno-cortical Carcinoma. Cancer Cell 29, 723–736. 10.1016/j.ccell.2016.04.002.

